# Cryo-EM structure of the human cardiac myosin filament

**DOI:** 10.1101/2023.04.11.536274

**Authors:** Debabrata Dutta, Vu Nguyen, Kenneth S. Campbell, Raúl Padrón, Roger Craig

## Abstract

Pumping of the heart is powered by filaments of the motor protein myosin, which pull on actin filaments to generate cardiac contraction. In addition to myosin, the filaments contain cardiac myosin-binding protein C (cMyBP-C), which modulates contractility in response to physiological stimuli, and titin, which functions as a scaffold for filament assembly^1^. Myosin, cMyBP-C and titin are all subject to mutation, which can lead to heart failure. Despite the central importance of cardiac myosin filaments to life, their molecular structure has remained a mystery for 60 years^2^. Here, we have solved the structure of the main (cMyBP-C-containing) region of the human cardiac filament to 6 Å resolution by cryo-EM. The reconstruction reveals the architecture of titin and cMyBP-C for the first time, and shows how myosin’s motor domains (heads) form 3 different types of motif (providing functional flexibility), which interact with each other and with specific domains of titin and cMyBP-C to dictate filament architecture and regulate function. A novel packing of myosin tails in the filament backbone is also resolved. The structure suggests how cMyBP-C helps generate the cardiac super-relaxed state^3^, how titin and cMyBP-C may contribute to length-dependent activation^4^, and how mutations in myosin and cMyBP-C might disrupt interactions, causing disease^5, 6^. A similar structure is likely in vertebrate skeletal myosin filaments. The reconstruction resolves past uncertainties, and integrates previous data on cardiac muscle structure and function. It provides a new paradigm for interpreting structural, physiological and clinical observations, and for the design of potential therapeutic drugs.

## Introduction

The heartbeat results from interaction between myosin and actin, present in the heart as organized arrays of protein filaments that slide past each other to produce contraction. After each beat (systole), the heart enters a relaxed state (diastole), leading to filling of the chambers with blood before the next contraction, a process that continues, without break, for the entire lifespan. Contraction is modulated by modifications to the myosin and actin filaments brought about by external stimuli.

Cardiac myosin (“thick”) filaments are bipolar polymers of myosin II^7^, whose α-helical tails form the filament backbone, and paired heads, lie at the surface^1, 2, 8, 9^. Each head consists of a motor domain (MD), essential light chain (ELC) and regulatory light chain (RLC). Three-fold symmetric “crowns” of heads form quasi-helices with a repeat of 430 Å **(Fig. 1a)**. Two myosin-binding proteins are also present. cMyBP-C binds at nine 430 Å-spaced sites in the middle 1/3 of each half filament, forming the C-zone, while strands of the giant protein titin run along each half-filament, from the tip to the filament center **(Fig. 1a)**. Titin and cMyBP-C consist of strings of globular, immunoglobulin-like (Ig) and fibronectin-type-III-like (Fn), domains that interact with myosin heads and tails to define thick filament architecture (titin) and regulate myosin function (cMyBP-C)^1^.

**Figure 1.**
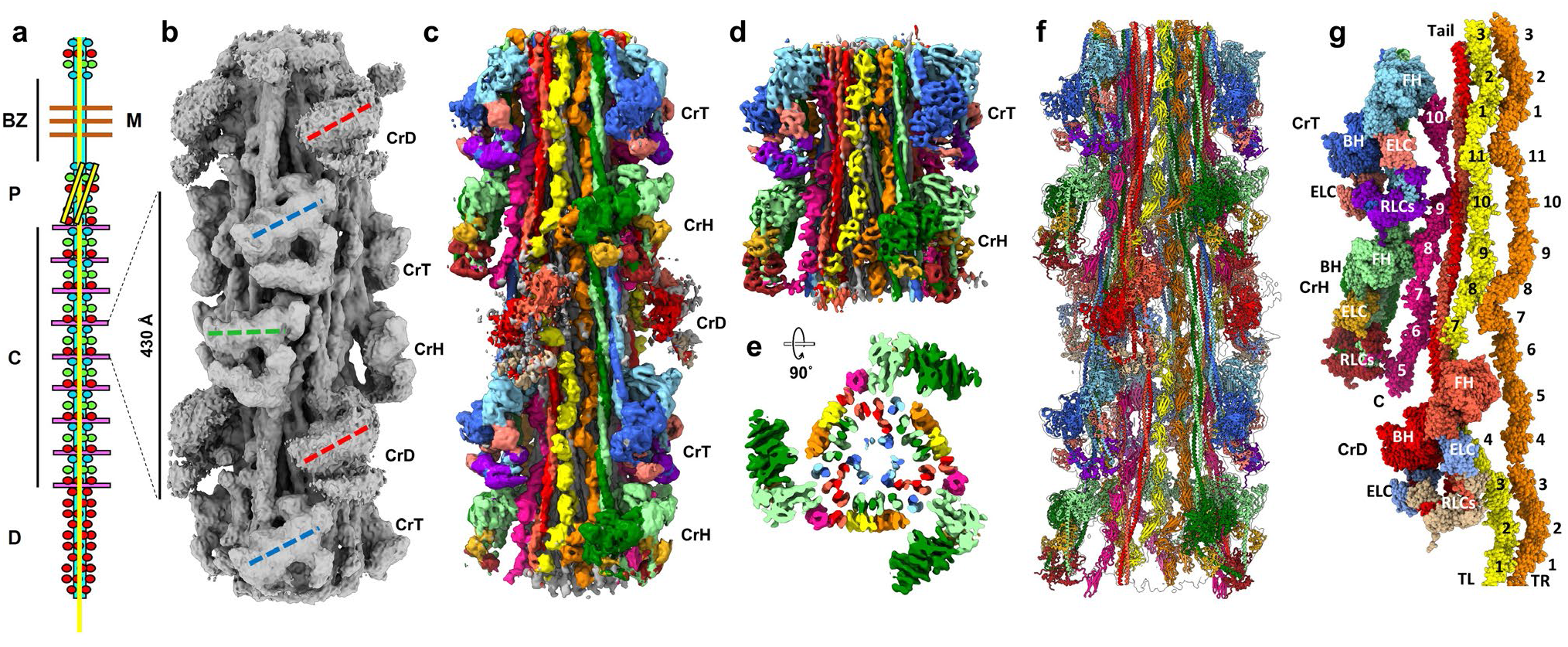
Single-particle 3D reconstruction of C-zone reveals organization of titin, cMyBP-C, and myosin heads and tails. **a,** Cartoon of half thick filament, defining P-, C- and D-zones, bare zone (BZ) with central M-line, quasi-helical arrangement (angled yellow lines) of myosin heads (colored spheres), with 430 Å repeat, and locations of cMyBP-C (pink) and titin (yellow). **b,** 3D reconstruction of 5 crowns of heads, longitudinal surface view, showing IHMs forming 3 types of crown (horizontal, CrH, green; tilted, CrT, blue; and disordered, CrD, red). Oriented with M-line at top. **c,** 5-crown map colored to identify myosin heavy chains (CrH, light/dark green; CrT, light/dark blue; CrD, light/dark red), coiled-coil myosin tails, and longitudinally arranged cMyBP-C molecules (pink) and 2 titin strands (yellow and orange); other colors in heads are light chains. **d,** Two-crown map (CrH and CrT) yields higher resolution, revealing α-helices in heads and tails. **e,** Two-crown map viewed transversely at the level of CrH, showing azimuthal and radial positions of components, and demonstrating resolution of myosin tails into individual α-helices (dark and light color within each tail). **f,** Five-crown reconstruction (silhouette) fitted with atomic models of myosin, cMyBP-C, and titin (ribbon depiction). **g,** Atomic model of portions of two sectors of reconstruction (defined as in **ED Fig. 2**), oriented to optimize visibility of all components. C (cMyBP-C) and TR, TL (right and left titins), with domains numbered; BH, FH, blocked and free heads; RLC, regulatory light chain; ELC, essential light chain.

During diastole, myosin filaments are switched off by intramolecular interaction of their paired heads (blocked, BH and free, FH^10^), forming interacting-heads motifs (IHMs^11, 12^), found across the animal kingdom^13^. Sequestration of myosin heads by the IHM appears to underlie the super-relaxed (SRX) state of myosin^3^ (most highly developed in the C-zone^14, 15^), in which ATP turnover is highly inhibited, conserving energy, and providing a reserve of myosin heads for increased physiological demand^16^. cMyBP-C and the myosin RLC are both phosphorylated in response to stimulation, leading to enhanced contractility.

Myosin, cMyBP-C, and titin all exhibit mutations that can cause dilated or hypertrophic cardiomyopathy (DCM or HCM), leading to heart failure^1^. HCM is typically associated with mutations in myosin and cMyBP-C, while titin is the most commonly-mutated protein in genetically-associated DCM. Myosin mutations frequently occur in the IHM interfaces, disrupting head-head interactions and the SRX state, and leading to the observed hypercontractile phenotype^5, 6^. Mavacamten, recently approved by the FDA for treatment of HCM, stabilizes the IHM and SRX^17^.

Despite the essential importance of the myosin filament to normal cardiac function and disease, its molecular structure remains poorly understood after 6 decades of research. We have determined the three-dimensional (3D) structure of the human ventricular filament C-zone at 6 Å resolution, revealing in molecular detail the complex organization of myosin heads, tails, titin and cMyBP-C. The structure provides solutions to past controversies and a new paradigm for interpreting studies of cardiac structure and function.

## Results and Discussion

### Structure of C-zone at 6 Å resolution

We carried out a single-particle 3D reconstruction of the 430 Å repeat unit of the C-zone of native, frozen-hydrated, human ventricular myosin filaments (EM Data Bank codes EMD-XXXXX, EMD-XXXXX, EMD-XXXXX). Global resolution was ∼6 Å (6-fold better than previous negative stain reconstructions^18, 19^, and locally as high as 4.5 Å; **Extended Data (ED) Fig. 1**). The interior architecture of the filament was revealed for the first time. The 3-fold symmetric reconstruction comprised 9 pairs of myosin heads on the filament periphery, segments of ∼33 myosin tails in the backbone, with 3 cMyBP-Cs, and six 11-domain titin strands on its surface: a total of 123 polypeptide chains and a MW of ∼ 5.9 MDa (Protein Data Bank (PDB) code XXXX; **Fig. 1b-g**). We first identify these different components, then discuss the interactions between them that create the complex and underlie its function.

#### Myosin heads

Myosin head pairs were clearly recognized as IHMs forming three distinct “crowns”: CrH, CrT, CrD, named for their appearances (**Fig. 1b; ED Fig. 2** for abbreviations), arranged in triplets, in a 3-start quasi-helical arrangement^18, 19^. CrH and CrT IHMs had similar conformations, but different orientations: CrH, *Horizontal*; CrT, *Tilted* (20° counterclockwise compared with CrH). IHMs in the third crown were similarly tilted, but weaker and noisier, suggesting significant head mobility (CrD, *Disordered*; **Fig. 1b, ED Fig. 1**). There was little azimuthal rotation from crown H to T (16°), but a larger rotation to CrD (32°), then 72° to the next CrH; there were small differences in axial spacing between crowns. Preliminary docking showed that all 3 crowns fitted well with a tarantula-based “canonical” IHM^20^ (PDB 5tby), supporting the hypothesis that the overall structure of the IHM is conserved across animal evolution^13^. However, the different tilts and stabilities of IHMs in the 3 crowns, and their distinct molecular environments (due to interactions with other components, described later), suggest diverse functional capabilities—contrasting with invertebrates, where all IHMs are in equivalent environments^11, 21^. Final docking was carried out by flexible fitting of X-ray crystallographic structures for the heads and tails to generate an atomic model (PDB XXXX; **Fig. 2**) that was used in subsequent analysis (see Methods). Strikingly, IHMs were at much greater distance (40 Å) from the filament axis than thought from previous negative stain reconstructions^18, 19^ (**ED Fig. 3**), as also inferred from recent X-ray modeling of intact muscle^22^. Agreement of X-ray and cryo-EM data suggests that the reconstruction closely resembles the filament structure in situ, highlighting its relevance to in vivo cardiac physiology.

**Figure 2.**
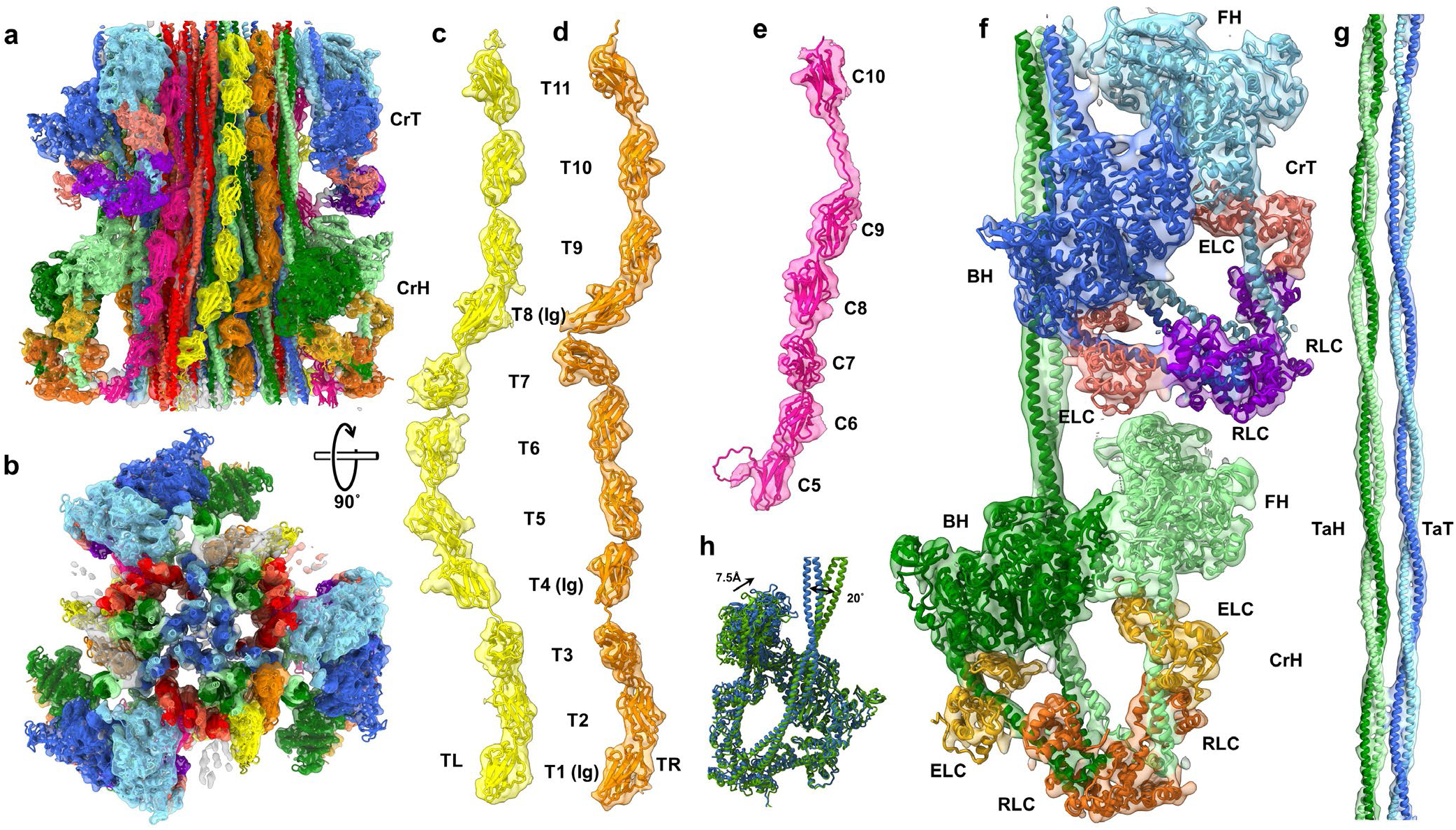
Model building and fitting explains all observed densities in the cryo-EM map. **a,** Refined atomic model (see Methods) fitted into translucent map of the two stable crowns (CrH, CrT, **Fig. 1d**); the model also shows titins, TL (yellow) and TR (orange), and cMyBP-C (C5-C10, pink). **b,** Same as **a**, rotated 90° (viewing towards Z-line). CrT tails (blue) form filament core, CrH tails (green) are distributed between surface and core, and CrD tails (red) are at filament surface, forming docking station for cMyBP-C (pink) (see **Fig. 5, ED Fig. 8**). **c-g,** Map densities of individual components (“split-maps”) segmented from full map, with corresponding models, to show detailed fitting. **c, d,** TL, TR, with domains numbered, and indicating the 3 Ig domains (others are Fn). TL and TR conformations differ mostly in middle domains T4-T7 (Ig-Fn-Fn-Fn). T1-T3 (Ig-Fn-Fn) and T8-T11 (Ig-Fn-Fn-Fn) are parallel to each other in corresponding regions of TL and TR **(ED Fig. 6)**. **e,** cMyBP-C domains C5-C10. Note extended C9-C10 linker predicted by AlphaFold **(ED Fig. 5)** is seen directly in map. **f,** CrH and CrT IHMs. **g,** CrH and CrT tails, TaH and TaT. **h,** CrT and CrH IHMs appear very similar when superimposed, except for a 7.5 Å shift between the two FHs, and 20° difference in angle between S2’s after they leave the BH.

**Figure 3.**
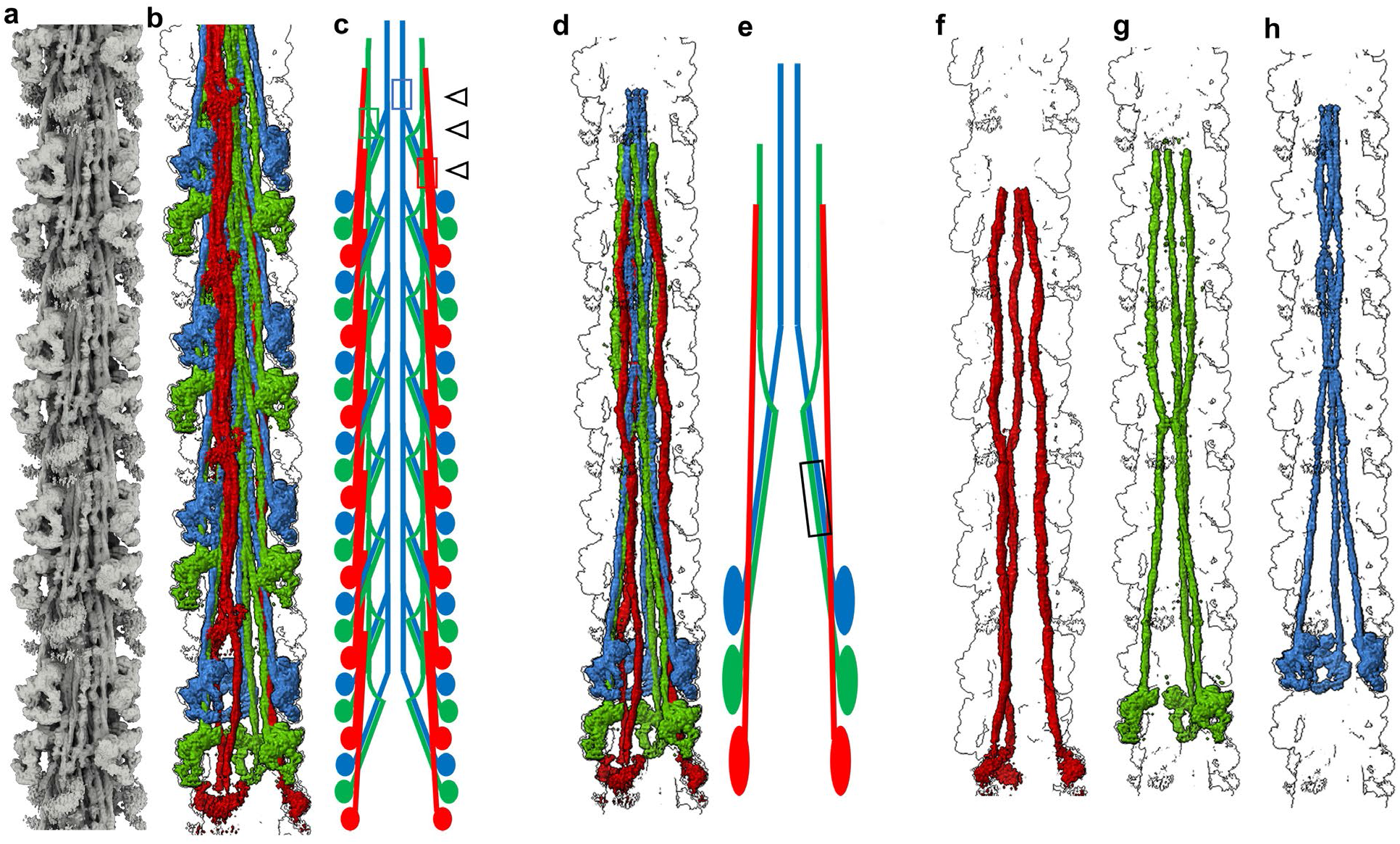
Myosin tails form an interconnected network in the filament backbone. **a,** Density map extended to 14 crowns to include full length myosin tails, revealing the network of interactions in the filament C-zone. **b,** Density map of multiple 430-Å repeats, colored to separate CrH (green), CrT (blue) and CrD (red) tails, highlighting their organization in the tail network. **c,** Schematic representation of tails from two of the three sectors of the filament, based on **(b)**. The most common interactions involve tails of myosins staggered by 3 crowns (430 Å; **ED Fig. 4a, e**). 430-Å-staggered CrT tails (blue) interact with each other in the distal half of LMM (representative interaction shown by blue box and black open arrow), forming the cylindrical filament core (**Fig. 2b, ED Fig. 4a, b**). Groups of 430-Å-staggered CrD tails (red) form a flat sheet (red box and black open arrow), on which cMyBP-C docks (**Fig. 5, ED Figs. 4a, b, 8**). CrH tails (green) also mostly interact with a 430-Å stagger (green box and black open arrow). **d-e,** Density map of myosin molecules of a single 430-Å repeat **(d)**, with schematic diagram **(e)**, showing mainly the single crown (141 Å) staggered S2-S2 interaction between CrH and CrT tails (black box). Other staggers between tails (5 and 7 crowns) are shown in **ED Fig. 4f**. **f-h.** Density map showing how myosin tails originating from the three different crowns in a 430-Å repeat follow three different trajectories: **f,** CrD; **g,** CrH; and **h,** CrT. This unique pattern differs fundamentally from invertebrate filaments, where all myosin tails are equivalent^21^.

#### Myosin tails

To examine the filament backbone, we extended the reconstruction to 14 crowns (**Fig. 3a;** see Methods), sufficient to reveal full-length myosin molecules, and focused on their organization in one of the 3 equivalent 120° sectors **(ED Fig. 2c)**. Myosin tails, well-resolved from each other, and measuring ∼1590 Å in length (consistent with sequence analysis^23^), were unambiguous, from N- to C-terminus, due to the characteristic left-handed super-coiling of their paired α-helices **(Figs. 1-3, ED Fig. 4).** Crossovers of the two helices were typically ∼75 Å apart, but longer in the regions containing the 4 skip residues (T1188, E1385, E1582, G1807), where the coiled-coil α-helices became more parallel or sharply bent, consistent with distortion of coiled-coil structure by skip residues **(ED Fig. 4h)**^23, 24^.

**Figure 4.**
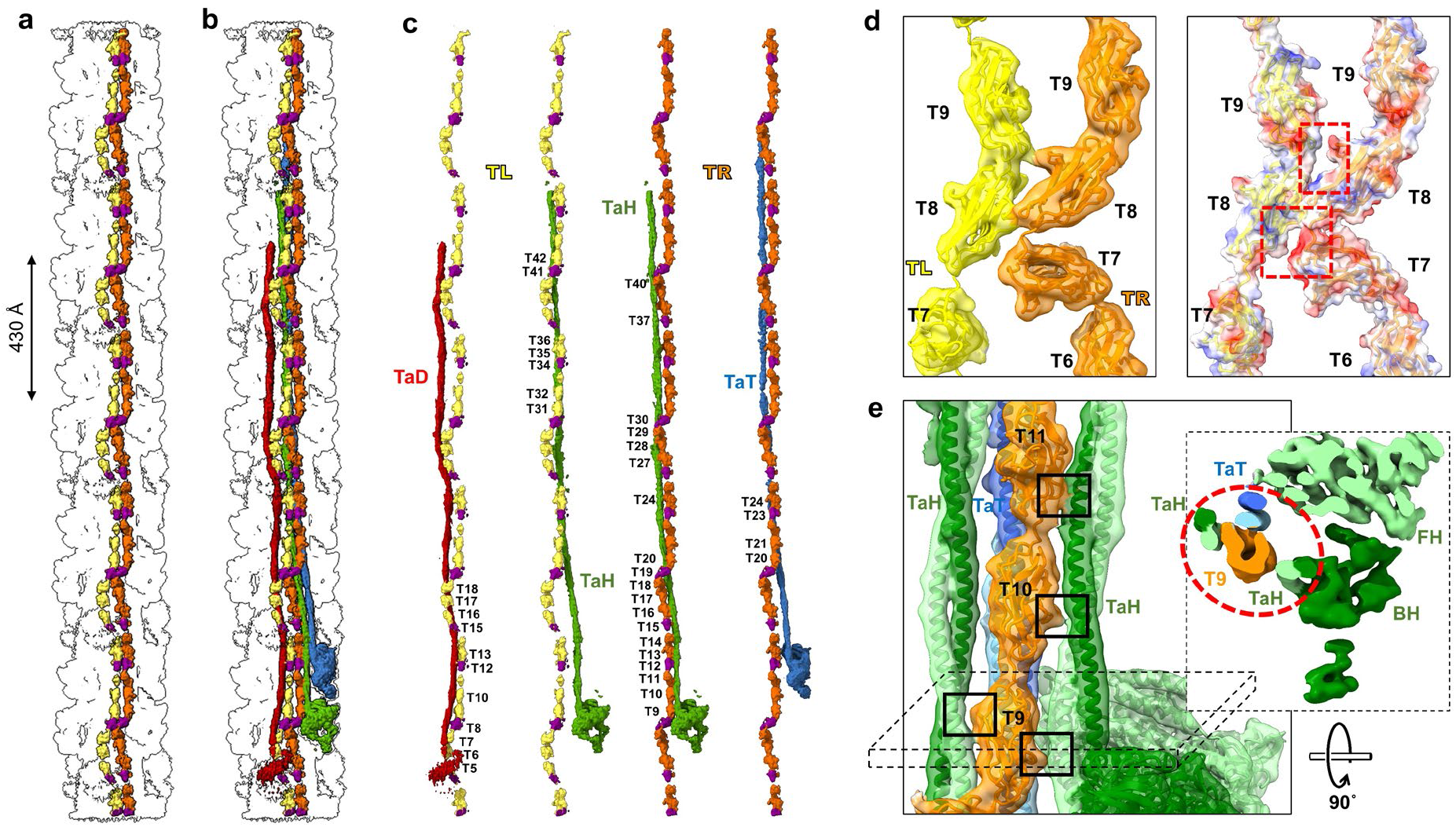
Titin functions as a template for myosin organization. **a,** Density map showing one pair of titin strands (TR, orange Fn domains, purple Ig domains; TL, yellow Fn, purple Ig). The two titins run nearly parallel except in the middle region (Ig-Fn-Fn-Fn) of each super-repeat (**Fig. 2c, d, ED Fig. 6**). **b,** Same as **a**, but including one 430-Å repeat of myosin molecules (CrH, green; CrT, blue; CrD, red), to show organization of tails with respect to titins. **c,** Titins can form extensive interactions with myosin tails in a sector **(ED Fig. 2d)**, creating and strengthening the tail network. Interactions of TL and TR with individual tails from CrH, CrT and CrD (TaH, TaT and TaD, respectively) are shown. Numbers indicate the individual titin domains involved. TaH interacts with both TL and TR: the first half (S2) extensively with TR, the second half (LMM) with TL. TaT interacts only (and minimally) with TR, mainly in its S2 region, and then travels towards the filament core **(Fig. 3)**, with no further interaction with either titin. TaD (mainly S2) interacts with TL, mostly in the mid-region (Ig-Fn-Fn-Fn) of two super-repeats (T5-T8 and T15-T18), but not with TR. **d,** TL and TR from adjacent sectors interact with each other to create a complete, 3-sector, filament. The main interaction, at T7/T8, is electrostatic (left, atomic model fitted into EM map; right, surface charge interaction). **e,** Representative example of titin-tail interface, showing TR (T9-T11) interaction with two TaH’s from different crowns and a TaT. Fitting of tail and titin atomic models into the map suggests β-β hairpin loops in Ig and Fn domains mostly form the titin-tail interface (left, black boxes). A single titin domain can interact with as many as 3 myosin tails originating from different crowns (right, inset). See also **ED Fig. 6**.

The tails showed no sign of association into subfilaments^25^, nor did they follow a “molecular crystal” arrangement of the type proposed by Squire^26^, and demonstrated in invertebrate filaments^21^, where all tails have equivalent environments: vertebrate filaments have a unique tail organization, not predicted by any previous model. Tails started (as S2) at the junction of the heads of each IHM, but then followed 3 quite different paths to their C-termini, depending on their crown of origin **(Fig. 3, ED Fig. 4).** CrH tails dipped initially toward the filament backbone, then reverted toward the surface (near skip 2, ∼755 Å from its origin). CrT tails started similarly, in contact with CrH tails, but then stayed at low radius, forming the inner cylindrical core of the backbone **(Fig. 3, ED Fig. 4)**. CrD tails were completely separate, always near the filament surface, where they made many interactions with cMyBP-C and titin. Their S2 regions had a flattened appearance (red/orange tails, **Fig. 1c, d**), suggesting vibrational motion parallel to the plane of flattening, consistent with the mobility of the CrD heads. The different courses of the 3 types of tail thus correlate with the different crown structures from which they originate, possibly contributing to functional specialization of these crowns, enabling a flexible response to physiological demand. Such specialization represents an evolutionary advance over the (more primitive) invertebrates, with their simple arrangement of myosin, in which all molecules are structurally equivalent, in a strictly helical organization, which would produce a more stereotyped response^11, 21^. The unexpectedly complex, non-equivalent arrangement of tails in the native vertebrate filament suggests caution in interpreting structure/function in synthetic filaments made from purified myosin.

#### Titin and cMyBP-C

In association with myosin, strings of ellipsoidal subunits, ∼40 Å long, ran along the backbone surface, suggestive of the linear arrangement of globular, ∼10-kDa, Fn and Ig domains in titin and cMyBP-C^27–31^ **(Fig. 1)**. In each 120° sector (**ED Fig. 2c, d**), two 11-domain strands ran parallel to and in register with each other, consistent with the two strands of titin’s 11-domain 430-Å C-zone super-repeat (Ig-Fn-Fn-Ig-Fn-Fn-Fn-Ig-Fn-Fn-Fn) and its stoichiometry of 6 molecules her half filament^32^; we refer to these as left and right titins (TL and TR) **(Figs. 1, 2, ED Fig. 2)**. The two strands do not appear to interact, except at two intermolecular contacts, Ig domains T1 and T8 **(Fig. 4d)**, marking the junction between sectors. Their flexible path, kinked at each Ig domain, is consistent with short, pliant linkers between domains^28^, but there is no evidence for extended dimerization or helical twisting of strands about each other^28, 33^ **(Figs. 1, 2, ED Fig. 6)**. We also observed a single strand with 6 subunits, consistent with the C-terminal half (C5-C10) of cMyBP-C, which includes domains C8-C10, thought to anchor cMyBP-C to the filament backbone **(Figs. 1, 2)**^30^. We validated these interpretations, and uniquely identified each domain of cMyBP-C and titin, using AlphaFold^34^ (**ED Fig. 5**). Our interpretation, based only on structure, predicts that both titin and cMyBP-C have their C-termini closer to the M-line, directly verifying previous conclusions from sequence analysis and antibody labeling^35, 36^.

**Figure 5.**
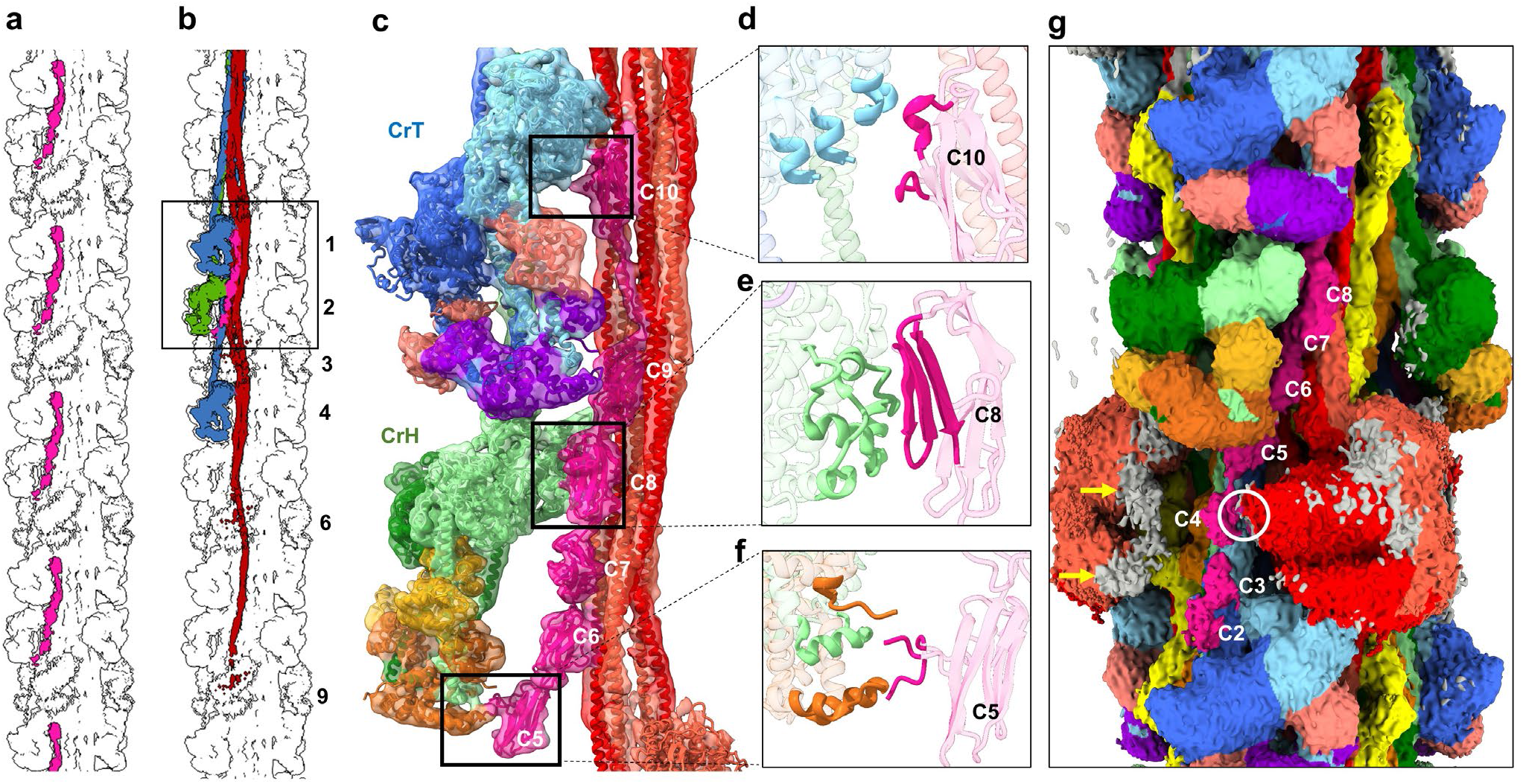
cMyBP-C interacts with myosin tails and heads. **a, b,** Overview of 13-crown map showing **(a)** positions of cMyBP-C (pink) at 430-Å intervals in one sector of the 3-fold symmetric filament, and **(b**) interaction of cMyBP-C with five myosin molecules (box), including CrH and CrT heads and a sheet of tails from three 430-Å levels (3, 6, and 9) of CrD. **c,** Zoom of map with fitted atomic model shows interaction of domains C6-C10 with CrD tails (see **ED Fig. 8** for details), C10 with CrT-FH-MD **(d)**, C8 with CrH-FH-MD **(e)**, and 28-amino-acid insert in C5 with CrH-FH-RLC **(f). g,** Low contour cutoff surface view reveals significant density for domains C2-C4, not observed at higher contour cutoff **(Fig. 1c)**; C2 appears to bind to CrT BH at bottom and C4 to CrD BH (white circle). Gray on CrD FH map (orange) lies outside CrD atomic model, possibly representing weakly (transiently) bound C0-M domains of cMyBP-C (yellow arrows). Analysis of our preparations using Pro-Q Diamond/SYPRO Ruby-stained gels suggests a basal level of cMyBP-C phosphorylation, which may contribute to the weakness of C0-M visibility^43^.

**Figure 6.**
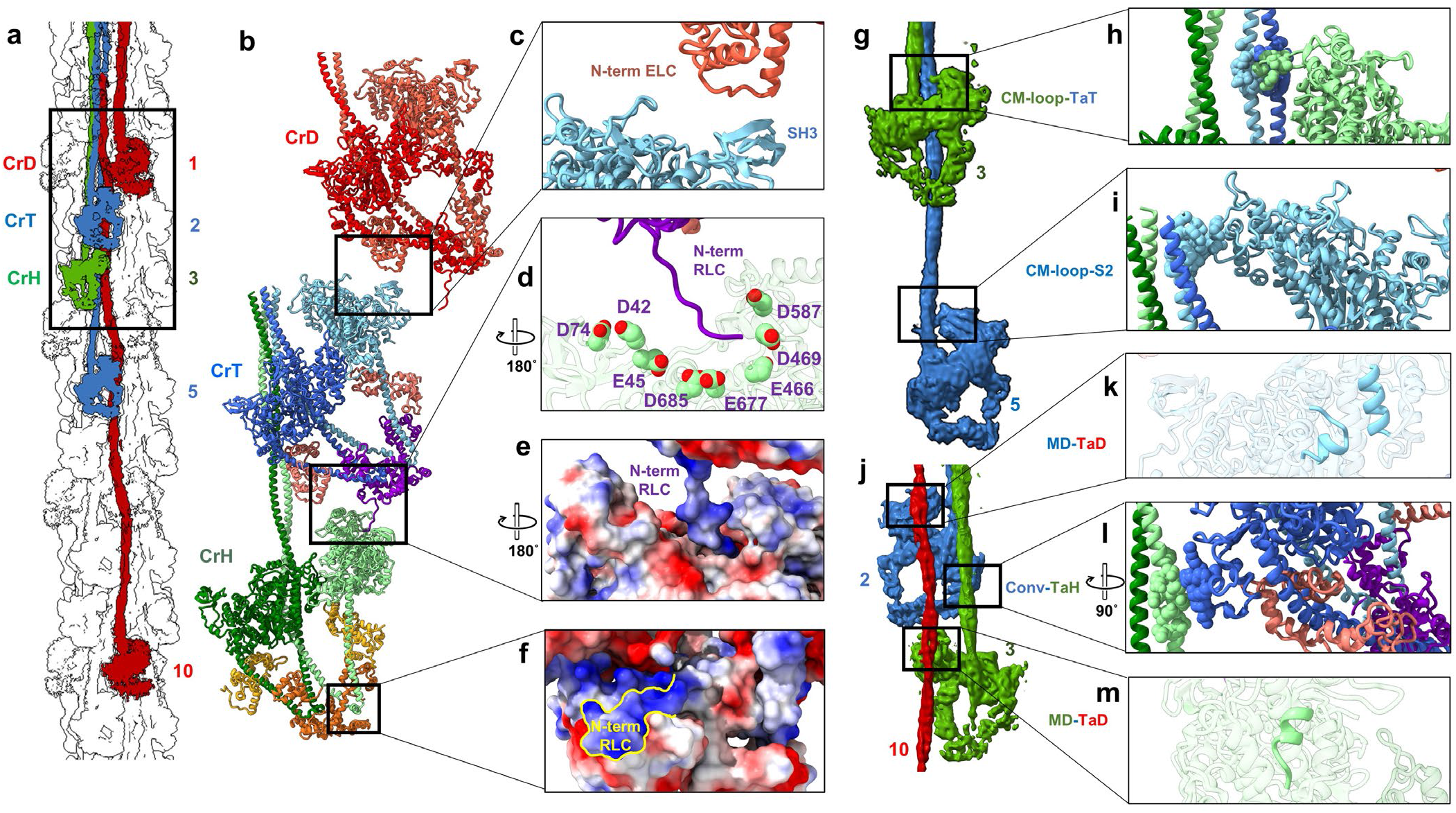
IHM interactions stabilize the relaxed state. **a.** Density map showing one CrH-CrT-CrD triplet, and CrT and CrD IHMs from two other triplets. **b.** Atomic models of IHMs in three crowns, showing possible inter-crown contacts. **c**. N-terminal ELC of CrD-BH is close to SH3 domain of CrT FH MD. Although the N-terminal 38 amino acids of the ELC are missing/disordered from the model (due to absence from PDB 5n69, used to create it), these could easily contact the CrT SH3 domain. **d.** AlphaFold-based model of positively charged N-terminal region of CrT-BH RLC shows possible contacts with negatively charged cavity of MD of CrH-FH (mostly Asp and Glu residues). This could help immobilize both CrT and CrH IHMs (enhancing SRX), and phosphorylation on Ser15 may weaken this contact^61^. **e.** Electrostatic surface representation of **d** (blue, positive; red, negative). **f,** The positively charged N-terminal extension (NTE) of the FH RLC (yellow outline) docks onto a negatively charged region of the BH RLC, establishing an RLC-RLC interaction. On activation of cMLCK, Ser15 is phosphorylated, which is thought to elongate the NTE^61^, in turn disrupting the RLC-RLC interaction and aiding release of the CrH FH. **g-m,** IHMs also make contact with different tails. **g, h, i.** The CrH FH CM-loop interacts with TaT (S2) from the crown below **(h)**, while the CrT CM-loop contacts its own S2 **(i)**. **j-m,** CrT and CrH FH interact with the same TaD from a lower level CrD through α-helices in their MDs **(k, m)**. **l,** The CrT-BH converter has possible contact with CrH S2.

In summary, the reconstruction fully accounts for the major known components in the C-zone of the vertebrate thick filament, and for the first time reveals their organization at high resolution.

### Interactions between components

Specific interactions between components are central to creating a biomolecular complex and determining its function. Myosin, titin, and cMyBP-C all have multiple domains and extended structures, making numerous interactions possible. Many have been inferred from studies over 6 decades. Our reconstruction reveals the major interactions in the native filament **(ED Fig. 2)**, including numerous contacts of myosin heads with heads, tails with tails, titin with tails, and cMyBP-C with heads and tails—but, unexpectedly, no interaction of titin with myosin heads or with cMyBP-C (**Figs. 1, 2, ED Fig. 2**). We first describe the tail-tail and titin-tail interactions that determine filament architecture, then the head–cMyBP-C, tail-cMyBP-C and head-head interactions that define its function.

#### Myosin tails form an interacting network

Interaction between tails is well-known to underlie myosin filament formation, through charge-charge attraction. Muscle X-ray patterns^37^, EM of myosin tail paracrystals^2, 38^, and sequence analysis^23^ suggest important tail staggers of ∼143 Å and 430 Å, arising from a periodic charge repeat every 28 residues (42 Å) along each tail: attraction is predicted between tails displaced by ∼(*n* + ½) x 28 residues (*n* integral)^23^. The reconstruction shows that, within one triplet of IHMs (CrH-CrT-CrD), interacting CrH and CrT S2s are staggered by ∼140.6 Å, directly revealing the predicted charge attraction for a 95-96 residue (141-143 Å) stagger of S2s^23^ **(ED Figs. 4g, 7a)**. But the full network of tail interactions occurs only when tails from axially neighboring triplets contact each other **(Fig. 3a-c, ED Figs. 4a-f)**. A key example is the merging of the light meromyosin (LMM) regions of three CrT tails from one triplet with three from the next, creating the cylindrical core described earlier **(Fig. 3, ED Fig. 4a, b, e)**. This type of interaction, with a 430 Å stagger between tails of the same type, is the most common in the reconstruction, occurring also with CrH and CrD tails **(ED Figs. 4a** (neighboring tails with crown number differing by 3); **4e**). It is made possible because crowns of the same type, 430 Å apart, have the same azimuthal and radial positions, placing their tails close enough for mutual electrostatic interaction^23^ **(ED Fig. 4g)**. Interactions between tails from different crown types (CrH-CrT, CrT-CrD, CrD-CrH) also occur, less frequently, when staggered by 5 or 7 crowns (715 or 1001 Å; **ED Fig. 4d,f)**, as predicted by sequence analysis^23^. The 3-, 5- and 7-crown staggers between tails all involve the distal region of LMM, which includes the C-terminal assembly competence domain (ACD, residues 1871-1899; **SI Table 1b**), consistent with its essential role in filament formation^39^ **(ED Figs. 4d-f)**. These varied tail interactions, differing depending on crown of origin, and localized to specific points along the tails, contrast with the simpler, helical invertebrate backbone, where all myosin molecules are equivalent, and tails interact along most of their length^21^.

#### Titin dictates assembly of tails

The above tail network is only possible if myosin molecules are placed at particular positions axially, azimuthally, and radially. Our data suggest that this occurs through their positioning by titin, with its 430 Å super-repeat functioning as a C-zone template^27^. The two titin strands in each sector show numerous interactions with myosin tails, mostly CrH and CrD, and a few with CrT **(Fig. 4a-c).** Both Ig and Fn titin domains are involved. CrH S2 interacts with all 11 domains of TR via attraction of the ∼40-Å–spaced +/− charge repeat on S2^23^ to complementary charges on the ∼40-Å–spaced titin domains **(ED Fig. 7b)**. LMM of the CrH tail also interacts with TR, and to a similar extent with TL **(Fig. 4c)**. Titin’s positioning of CrH myosins 430 Å apart is reinforced by the 430-Å–staggered tail-tail charge interactions described above^23^ **(ED Fig. 4g)**. In contrast to CrH, CrT tails interact little with titin, being positioned instead by the ∼140-Å-staggered electrostatic attraction of CrT S2 to CrH S2 described above **(Fig. 3e, ED Figs. 4g, 7a)**. The extensive interaction of CrH (and to a lesser extent CrT) tails with TR occupies many TR binding sites; in fact, an individual titin domain may bind as many as 3 tails around its surface **(Fig. 4e)**. This may explain the need for a second titin (TL) in each sector, to position CrD tails (**Fig. 4c)**. This interaction, with S2 of CrD **(Fig. 4c)**, occurs in the middle sections of two TL super-repeats (T4-T7), which are rotated ∼90° relative to TR (between kinks at T3/T4 and T7/T8), placing this region at higher radius **(ED Fig. 6a, b)**, and with different titin binding sites exposed. This may explain CrD tails forming the outer shell of the backbone **(ED Fig. 4a)**, their axial displacement from CrT, and the greater distance of CrD heads from the filament surface. As CrH and CrT tails interact with the same titin (TR), CrH is close to CrT azimuthally. In contrast, the interaction of CrD with TL, where TL and TR in the same titin pair are ∼48 Å apart, explains the large azimuthal rotation (∼72°) from CrD to CrH of the next triplet **(ED Fig. 6d)**.

Compared to the extensive interaction of titin with myosin tails, we see no interaction with myosin heads. This supports some in vitro studies of purified proteins^40^, but contradicts others, which concluded that titin binds to myosin mainly through the heads^41^. The reason for the discrepancy between previous in vitro data is unclear, but comparison with our observations suggests caution in using such data to decipher native structure.

In summary, we see the detailed course of all 11 domains of both titin strands, which position the myosin tails in a unique network arrangement, whose interactions are reinforced by electrostatic tail-tail interactions first predicted 40 years ago.

#### cMyBP-C binds tails, heads but not titin

cMyBP-C^30^ is a modulator of myosin activity in the C-zone, where it is thought to enhance myosin’s SRX state^14, 15, 42^, moderated by phosphorylation of the M-domain^43^. With no previous high-resolution information on its organization, it has remained a mystery how cMyBP-C, with a ratio of only one to every 3 myosins in a 430-Å repeat, could affect the activity of multiple myosins^43^. The longitudinal orientation of cMyBP-C domains C5-C10 produces interactions with myosins from 6 different crowns **(Fig. 5)**: the FH MDs in crowns H and T and CrH FH RLC **(Fig. 5c-f)**, S2 from the next CrT away from the M-line **(Fig. 5b)**, and a complex of LMMs from 3 CrD crowns (**Fig. 5b, c, ED Fig. 8**). These interactions are consistent with in vitro studies demonstrating MyBP-C binding to S1, S2 and LMM^6, 30, 43^, but show that in the native filament, the interactions are confined to specific myosins. There is no evidence for a circumferential arrangement of cMyBP-C^30^, nor for interaction of a folded cMyBP-C conformation with the IHM^6^. Strikingly, the heads interacting with cMyBP-C are those of the two well-ordered crowns (specifically the FHs of CrH and CrT), suggesting that cMyBP-C contributes to this ordering, possibly stabilizing their IHM/SRX state and tethering them to the backbone.

In contrast with the longitudinal orientation of cMyBP-C’s C-terminal half, there is evidence that, in the sarcomere, its N-terminal domains extend perpendicular to the thick filament and bind to thin filaments^44^. In our preparation of thick filaments—lacking thin filaments—the N-terminal half of cMyBP-C is likely to be mobile, enabling it to bind weakly to the thick filament^43^. Our reconstruction supports this. At low contour cutoff, clear but weaker density is seen for domains C4, C3 and C2, loosely resting on S2 of the lower level CrT (**Fig. 5g**, TaT below C2-C4) and with possible contact of C4 with the CrD BH (**Fig. 5g**, white circle). While the most N-terminal domains (C0, C1 and M) are not visible as discrete subunits, they may contribute to additional weak density on the FH of CrD **(Fig. 5g,** yellow arrows). In vitro, cMyBP-C’s N-terminal domains C0-C2 can interact with both myosin and actin, implying possible switching between thick and thin filaments in different muscle states^43^. This is supported by X-ray diffraction of muscle at different sarcomere lengths^45^ suggesting that when thin filament binding sites are present (overlap zone of the sarcomere), N-terminal binding to actin produces the well-known 430-Å– spaced cMyBP-C stripes in the C-zone, but when absent (isolated filaments, sarcomeric H-zone), the N-terminal region is mobile. The weakness of thick filament/N-terminal binding in our reconstruction may be due to partial M-domain phosphorylation in our preparations **(Fig. 5)**, weakening binding to the filament^43^. Remarkably, the mobile region of cMyBP-C appears to be mainly at the level of the mobile—possibly disordered relaxed (DRX)—heads (CrD). These two regions might cooperatively guide each other to their interaction sites on the thin filament when present^46^. This location suggests that cMyBP-C’s most important functional region (C0-C2) would impact primarily CrD heads, for example upon M-domain phosphorylation, or loss of the N-terminal 29 kDa region through proteolysis during ischemia^43^.

Based on binding studies, it is widely assumed that titin positions cMyBP-C on the thick filament, through interaction of its C-terminus (C8-C10) with the first Ig domain of titin’s 11-domain super-repeat^47^. Surprisingly, we see no contact of cMyBP-C with titin: the two are clearly separated, by a CrD tail **(Figs. 1, ED Fig. 2a)**. Instead, cMyBP-C appears to dock onto the backbone by electrostatic binding of its C-terminal region to a sheet of 3 CrD tails lying side-by-side, originating from three consecutive 430 Å CrD repeats **(Fig. 5a-f; ED Fig. 8)**. Possibly, the C-terminal cMyBP-C interaction with titin seen in vitro occurs during filament assembly, but is lost in the mature filament; alternatively, titin determines cMyBP-C position indirectly, by dictating the positions of the CrD tails, as suggested by the reconstruction. Strikingly, C10 of cMyBP-C in the reconstruction aligns precisely with T1 of titin, while C8 and C9 extend over T9 and T10 of the previous super-repeat **(Fig. 1g)**. This agrees closely with conclusions from antibody labeling^36, 48^, providing strong support for our interpretation based on structure alone.

Division of the thick filament into C-and D-zones has been known for 40 years, but never satisfactorily explained^49^. It is usually attributed to cMyBP-C binding sites in titin’s C-zone 11-domain super-repeat that are absent from the 7-domain super-repeat of the D-zone^27, 43^. But this explanation fails if cMyBP-C does not bind to titin. The binding of cMyBP-C to the specific coalescence of CrD tails from three 430-Å-repeats **(ED Fig. 8)**, themselves positioned by titin’s 430-Å 11-domain super-repeat, would explain why cMyBP-C is confined to the C-zone, as this organization of tails would be absent from the 7-domain super-repeat of the D-zone. It would also explain why cMyBP-C is unexpectedly absent from the first two 430-Å titin repeats (those closest to the Z-line)^36^, starting instead ∼720 Å closer to the M-line^48^—because the full set of three 430-Å levels of CrD tails is required for cMyBP-C to bind.

In summary, cMyBP-C binds to myosin heads and tails, but not to titin. Its interactions, seen for the first time in the context of the native thick filament, suggest how it may modulate myosin activity, differently in different crowns, with fine-tuning by phosphorylation. Its longitudinal binding shows that in the absence of actin, as in our preparation, its N-terminal half (in addition to its C-terminal half) can bind to myosin, which has not been previously demonstrated structurally. Combined with evidence for actin binding, this provides strong support for models in which cMyBP-C’s N-terminal can switch binding partners, depending on actin-availability and muscle physiological state^43^.

#### IHM interactions modulate head activity

The reconstruction shows that, in addition to interacting with cMyBP-C, myosin heads interact with each other and with myosin tails. Within the IHM, the BH and FH MDs form a large interaction interface, and the BH interacts with S2, as in the canonical IHM of invertebrates^20^. These intramolecular interactions are thought to stabilize the SRX state of myosin heads in an inhibited (locked) conformation^50^. They can be studied at highest resolution by cryo-EM of isolated IHMs^51^, although we note that differences in CrH and CrT IHMs **(Fig. 2h)**, mean that isolated IHMs cannot precisely reproduce intramolecular interactions in all crowns of the native filament.

In addition to intramolecular interactions, contacts <u>between</u> IHMs appear likely along their quasi-helical tracks. These interactions, absent in studies of individual molecules^51^, may involve unstructured N-terminal regions of the ELCs and RLCs (**Fig. 6**). CrH, CrT and CrD form a triplet **(Fig. 6a, b)**, in which CrT and CrD are both tilted, such that the N-terminal residues of CrD BH ELC could contact the CrT FH SH3 domain below^19^ **(Fig. 6b,c)**—as in tarantula, where all IHMs are similarly tilted^11^. Intramolecular interaction of the N-terminal ELC with myosin SH3 has been previously suggested in vitro^52^. Our reconstruction indicates that in the thick filament *inter*-molecular interaction is possible. A key innovation in vertebrate thick filament evolution may be the 20° clockwise rotation of the CrH IHM relative to CrT (see *Myosin heads*, above; **Fig. 1b**). This brings CrH’s FH within range of CrT’s N-terminal BH RLC above^19^ **(Fig. 6b, d)**. Although no density is visible for the mobile RLC N-terminus, AlphaFold predicts that this positively charged region (1-MAPKKAKKR-9) could reach, and interact with, the cavity formed by 8 negatively charged amino acids (5 Asp and 3 Glu) of this FH **(Fig. 6d, e)**, possibly impacting Ser15 phosphorylation of the CrT BH RLC by affecting its accessibility to MLCK. This would enhance the plasticity of response of myosin activity over that in tarantula (where RLC interaction between crowns does not occur^11^)—a clear evolutionary advance. We also observe density for the N-terminal extension of the FH RLC (disordered in myosin head crystal structures), apparently stabilized by electrostatic interaction with the BH RLC **(Fig. 6f)**. Phosphorylation on Ser15 would be likely to break this interaction, releasing the CrH FH **(Fig. 6f).**

These contacts between CrH, CrT and CrD within a triplet suggest a physical communication pathway between crowns, which could contribute to cardiac function through local cooperative thick filament activation. Contact <u>between</u> triplets seems unlikely (CrH ∼ 50 Å from the preceding CrD), suggesting that triplets would function as individual cooperative units, responding independently to mechanical changes or signaling events. This would augment the versatility of response implied by the structural differences between crowns within a triplet.

There are also interactions between the heads and the tails. CrH and CrT FHs interact with the same CrD tail from 2 triplets below, via α-helices in their MDs **(Fig. 6j, k, m)**. The CM-loop of CrT-FH interacts with its own S2 **(Fig. 6g, i)**, while that of CrH-FH interacts with the tail of CrT below **(Fig. 6g, h; ED Fig. 9d).** The converter of CrT-BH also comes close to S2 of CrH below **(Fig. 6j, l)**. We suggest that these inter-IHM, IHM-tail (and also IHM-cMyBP-C) interactions together contribute to stabilization of SRX in the C-zone and may play a dominant role in filament activation (see *Physiological and evolutionary implications*, below).

### Structural consequences of mutations

Over 250 mutations in the myosin heavy chain, light chains, MyBP-C and titin are known to be pathogenic or likely pathogenic in humans, leading to HCM and DCM^1, 5^. Mapping these mutations on to the reconstruction suggests their likely structural impact. In summary, we confirm the clustering of mutations in the intramolecular interaction interfaces of the IHM suggested previously (based on a low-resolution homology map of the tarantula thick filament), which were proposed to weaken the IHM, impair relaxation and increase energy consumption, leading to HCM pathogenesis^5, 6^. A full analysis will be presented elsewhere. Here we highlight 4 representative examples of new or previously unexplained clusters of HCM mutations (mostly charge reversal or neutralization), whose locations in our map reveal their involvement in key, functionally important *inter*molecular interactions between myosin heads, tails, titin, and cMyBP-C. Strikingly, the different structures and interactions of the different crowns, and of the different heads (BH, FH) within them, means that a single mutation can have distinct effects at as many as 6 locations, or only affect one specific interaction **(ED Fig. 9)**. Mutations E924K and E930K in ring 2^53^ of CrD S2, at its interface with TL titin **(ED Fig. 9a)**, could weaken CrD binding to the filament and alter transmission of tension by titin to cMyBP-C, thus disturbing length-dependent activation (see next section). The same mutation (E924K) in a different (CrT) tail, at its interface with the CrH tail **(ED Fig. 9c)**, could have a quite different effect, destabilizing docking of the CrT IHM onto the filament, impairing its SRX state. Mutations in either cMyBP-C or myosin, at the C8/CrH FH interface **(ED Fig. 9b)**, could disrupt CrH FH stability and thus its SRX state. Finally, a classic mutation in the cardiomyopathy loop of the myosin head (R403W/G/L/Q) could have quite different effects depending on its location on the FH or BH^5^ and whether it is in CrH or CrT. On the CrT FH it could alter binding of the FH to its own S2, while on the CrH FH, binding to CrT S2 would be affected **(ED Fig. 9d)**. All of these disturbances are potential causes of HCM. We found no mutations in regions involving interactions between crowns.

### Physiological and evolutionary implications

Our reconstruction reveals the high-resolution structure and complex set of molecular interactions that underlie thick filament function in the human heart. A similar structure is likely in skeletal filaments, and across all vertebrates, based on their similar X-ray diffraction patterns^37, 54^, making this a new paradigm for vertebrate thick filament structure. The reconstruction provides many surprises and new understanding of the organization, energetics, function, and consequences of mutations in the cardiac thick filament. Three types of IHM characterize the C-zone, with different tail paths and different interactions. CrH and CrT are stabilized by cMyBP-C and likely represent SRX heads, while CrD IHMs are more mobile (DRX). The latter, along with mobile heads in the D-zone, may function as “sentinel” heads that detect thin filament activation and create initial tension in systole^14, 15, 55^. FHs are likely more stable than BHs, due to their greater number of interactions—the opposite of tarantula^56^. The flexible, finely-tuned response made possible by these different IHMs, with modulation by RLC and cMyBP-C phosphorylation, represents a major evolutionary advance over invertebrate thick filaments: lacking titin and cMyBP-C, and having a uniform set of tail and head interactions^11, 21^, invertebrate filaments would have a more stereotyped response for all molecules. Our structure suggests a plausible model for thick filament assembly in vivo in which titin is laid down first^27^ and recruits myosin II and finally MyBP-C into nascent filaments **(SI Discussion)**.

The heart must pump out what it takes in, and has developed a sophisticated mechanism for doing so. The Frank-Starling law is an auto-regulatory, beat-by-beat mechanism that adjusts the strength of contraction in systole to the amount of blood in the ventricles in the preceding diastole^4^. A fundamental concept of cardiology, it is equivalent at the cellular level to length-dependent activation (LDA): a longer cardiac sarcomere contracts more strongly than a shorter one. Numerous proposals have been made to explain LDA. Recent experiments suggest that activation of myosin heads (e.g. from SRX to DRX state), through increased stress on the thick filaments, transmitted passively via titin at longer sarcomere length, plays a key role^54, 57^. Our reconstruction reveals a simple molecular pathway involving titin, myosin and cMyBP-C, that could explain this mechanism **(ED Fig. 10)**. In this model, stretching of the sarcomere at end-diastole transmits tension via I-band titin to titin strands in the thick filament. TL and TR both exhibit spring-like kinks **(ED Fig. 6)**, that could be straightened under tension, causing CrD tails, attached to TL (**Fig. 4c**), to slide past each other, disrupting cMyBP-C’s main backbone binding site **(Fig. 5b, c, ED Fig. 8)**; this could disturb cMyBP-C’s stabilizing interactions with myosin heads, shifting the equilibrium from SRX towards DRX^57^. Additional CrH and CrT heads thereby made available for actin interaction would account for the enhanced systolic force that follows. Active force generated by myosin heads during systole, stretching the filament by up to 1.6%, may produce similar changes in CrD tails, cMyBP-C, myosin heads and titin, disrupting their interactions to explain thick filament activation by mechanosensing in both cardiac^58^ and skeletal muscles^59, 60^.

Our structure has important clinical relevance. Filaments were prepared in the presence of mavacamten, an FDA-approved drug used to treat HCM-based hypercontractility, by stabilizing the SRX of myosin heads^17^: we are therefore observing the structure of filaments as they might be in mavacamten-treated patients. We used mavacamten to enhance the resolution of our structure, but based on previous low-resolution reconstructions^18, 19^ and X-ray data^17^ it should not significantly alter the structure of the filament. Mavacamten has made possible new, high-precision insights into the structure and function of the native human cardiac filament and possible mechanisms of disease. Our structure unlocks the door to future planning of drug targets.

## Methods

### Research ethics for donated tissues

Tissue samples collected from the University of Kentucky (samples 5155D, 53018.4; 78952, 53132.3, and D0F54, 52902.3) were processed by the Gill Cardiovascular Biorepository. All samples came from the outer third of the left ventricular free wall. They were obtained from terminal organ donors whose hearts could not be used for transplantation for technical reasons (e.g. blood type mismatch). The local Organ Procurement Organization (Kentucky Organ Donor Affiliates, KODA) obtained informed consent from the legally authorized representatives, which allowed the hearts to be used for research if they could not be used as part of clinical care. All procedures were approved by Kentucky Organ Donor Affiliates and the University of Kentucky IRB.

### Myosin filament preparation

Cardiac ventricular tissue samples collected by technical staff in the operating room were immediately placed into cold saline, then taken to the lab for dissection into anatomic regions. They were flash-frozen in liquid nitrogen and stored in its vapor phase (see Blair et al. for further information^62^). Samples were sent on dry ice from University of Kentucky to University of Massachusetts Chan Medical School, and stored at -80°C. Myosin filaments were isolated from ventricular tissue under relaxing conditions. All steps were performed at room temperature (24°C). Chemicals were purchased from Sigma. A piece of frozen muscle (∼120-150 mg) was thawed by placing in relaxing solution (100 mM Na acetate, 8 mM MgCl2, 5 mM NaATP, 1 mM EGTA, 2 mM imidazole, 10 mM phosphocreatine, pH 6.8) with protease inhibitors (Leupeptin, Pepstatin and Aprotinin, 0.004 mg/mL each), and chemically skinned in relaxing solution containing 0.5% Triton X-100 for 30 min with agitation. The muscle was cut into small pieces and teased with forceps to make fine bundles^8^, which were further relaxed with fresh relaxing solution for 90 min. Relaxed bundles were treated with 0.25 mg/mL Porcine Pancreatic Elastase (Sigma, E0127, Type III)^8^ for 3 min, followed by addition of 1 mM PMSF for 10 min to stop the reaction. A filament homogenate was produced by gentle hand shaking for 3 min to release maximum numbers of myosin filaments, and the sample was clarified by centrifugation (3000 rpm, 3 min). Contaminating thin filaments in the supernatant were fragmented using the N-terminal half of gelsolin^63, 64^ (400 ug/mL), expressed in *E. coli* using a recombinant plasmid provided by Dr. Christine Cremo (University of Nevada). The resulting myosin filaments were pelleted by centrifugation (14000 rpm, 5 min), and dissolved and gently resuspended in 100 µL fresh relaxing solution containing 50 µM mavacamten. Filament quality was checked by negative staining^65^ with 1% w/v uranyl acetate before cryo-EM grid preparation.

#### Use of mavacamten

Mavacamten was used to stabilize the labile myosin head array and thus enhance the resolution of the reconstruction. X-ray diffraction suggests that it does this without significantly changing myosin head structure^17^. The overall similarity of myosin head conformation in our structure compared to negative stain reconstructions in the absence of mavacamten^18, 19^ supports this. 2D class averages of negatively stained cardiac filaments with and without mavacamten appear similar to each other (unpublished data), further implying no major change in structure. We avoided glutaraldehyde crosslinking, used in some cryo-EM studies to stabilize structure.

### Cryo-EM

3 µL of myosin filaments from the above 100 µL sample were applied to C-flat holey carbon grids (CF-1.2/1.3-3 Au, Protochips) glow-discharged for 45 sec at 30 mA using a PELCO easiGlow (Ted Pella). Grids were rear-side-blotted at 25°C and 95% relative humidity using a Leica EM-GP-2, and plunge-frozen in liquid ethane immediately after blotting. Grids were stored in liquid nitrogen until screening and data collection.

### Data acquisition

Grids were initially screened on a 200 keV Talos Arctica (Thermo Fisher Scientific) for ice thickness, filament quality, and filament distribution. Optimal grids were imaged at 300 keV on a Titan Krios (Thermo Fisher Scientific) equipped with Gatan K3 direct electron detector and Gatan GIF Quantum energy filter. Data collection (in counting mode) was performed in three different sessions, using filaments from 3 different hearts. A total of 56,489 movies were recorded (18,516, 18,872, and 19,101, respectively), using SerialEM V 4.0^66^, with 40 frames over 4s for a total electron dose of ∼60 *e^-^* Å^-2^. Nominal magnification was 105,000x, with a pixel size of 0.415 Å in super-resolution mode (physical pixel size of 0.83 Å at the specimen level), and a defocus range from -1 to -2 µm.

### Data processing

Images were processed using single particle methods; we avoided helical averaging, which would have lowered the resolution, as the filaments are only quasi-helical. Movies from the three Krios sessions were processed separately and the particles from all three sessions combined for final processing. Dose-fractionated movies were motion-corrected using MotionCor2^67^. Patch contrast transfer function (CTF) was used in CryoSPARC 3.3.2^68^ for CTF estimation. Initially, the reference-free filament-tracer module in CryoSPARC 3.3.2 was used to pick particles from all dose-fractionated micrographs, and the particles extracted using a box size of 1048 pixels (pixel size 0.83 Å), which included approximately 6 crowns of heads. The 1048-pixel box was Fourier-cropped to 800 pixels to produce a final pixel size of 1.0873 Å. Initial 2D-classification in CryoSPARC 3.3.2 (using 100 classes) resulted in multiple good classes, which were then used for template-based particle-picking. Next, iterative 2D classification was performed, resulting in 104928, 92790 and 149418 curated particles from the 3 Krios sessions, respectively. Particles were combined, and reference-free *ab initio* reconstructions with 4 classes were generated in CryoSPARC 3.3.2. These were further used as 3D models in heterogeneous refinement, using all selected particles from the 4 classes. The heterogeneous refinement resulted in the following particle numbers: class 0 (102581), class 1 (87325), class 2 (138288), and class 3 (18952). Particles from classes 0 and 2 were further refined using homogenous refinement, resulting in two final reconstructions, containing 102581 and 138288 particles, with estimated resolutions (gold-standard FSC 0.143 criterion) of 6.4 and 6.0 Å respectively. Local resolution was calculated in CryoSPARC 3.3.2. Maps were post-processed using DeepEMhancer^69^. Portions of the two final reconstructions containing the two stable crowns (CrH and CrT) were combined using the signal subtraction module in CryoSPARC 3.3.2, followed by 3D classification into 4 classes. 3D classification resulted in the following particle numbers: Class 0: 49227, Class 1: 82458, Class 2: 62747, and Class 3: 46436. Particles from classes 0, 2 and 3 were combined in a homogeneous refinement, resulting in a 2-Crown reconstruction with 158410 particles, showing more detailed secondary structure. A 13-crown extended map was generated by joining multiple 5-crown (CrH-CrT-CrD-CrH-CrT) maps, which were initially superimposed at the last two crowns (CrH-CrT) and first two crowns (CrH-CrT) of two 5-crown maps using the vop maximum command in ChimeraX.

### Model building and refinement

For myosin head model building, CrH was initially made from the crystal structure of the bovine β-cardiac myosin head (PDB 5n69), containing the motor domain and ELC region of the regulatory domain, which was rigid-body fitted to the CrH BH and FH of the 2-crown map, using *Fit in map* in ChimeraX. This was followed by molecular dynamics flexible fitting (MDFF, NAMDINATOR^70^) and manual building in COOT^71^. The S2 region was fitted using the crystal structure of the human β-myosin S2 fragment^53^ (PDB 2fxo). The RLC and its binding region on the lever arm were built using AlphaFold^34^ structures AF-P10916-F1-model_v3.pdb (the RLC) and the RLC-binding region of AF-P12883-F1-model_v1.pdb (the MYH7 heavy chain). The full IHM structure was finally refined using Phenix real space refinement^72^. To build CrT, the final CrH IHM was flexibly fitted (MDFF, NAMDINATOR) into the segmented density of CrT, followed by Phenix real space refinement. To construct the myosin tails, an initial model was built using AlphaFold Colab^34^ and then flexibly fitted into the map density starting from the head all the way to the end, in multiple iMODFIT normal mode refinement^73^/MDFF cycles, followed by Phenix refinement, as described above. AlphaFold database entry AF-Q14896-F1-model_v1.pdb was used as initial model for building cMyBP-C. Domains C5 to C10 from the model were fitted using MDFF and iMODFIT normal mode refinement **(Fig. 2, ED Fig. 5)**. For titins (TL and TR), titin C-zone 4 sequence was used to build the initial model, using AlphaFold Colab. The best fit to the map was obtained with domains T1-T3 (Ig-Fn-Fn) and T8-T11 (Ig-Fn-Fn-Fn), using ChimeraX Fit in Map. The middle domains T4-T7 (Ig-Fn-Fn-Fn) were fitted using flexible fitting (MDFF, NAMDINATOR). All parts were joined using COOT, and full-length TL and TR were refined using a final round of MDFF followed by Phenix real space refinement **(Fig. 2, ED Fig. 5)**. Iterative refinement was cycled between COOT and Phenix real space refinement for the full, 3-crown model. Validation was carried out using Molprobity^74^. Figures were made using UCSF ChimeraX^75^.

#### Model building of CrD

The full 3-crown atomic model (PDB XXXX) was built using a cryo-EM map with estimated overall resolution of ∼6 Å **(ED Fig. 2)**. Although CrD is mobile, with weaker density and lower local resolution, it too has an average IHM structure **(ED Fig. 2g)**. We therefore also included an IHM atomic model positioned to fit its overall low-contour density, showing where CrD is positioned (on average) with respect to the 2 stable crowns (CrH and CrT) in a triplet. This was carried out by rigid-body fitting of the CrT IHM atomic model into the CrD density using ChimeraX. Our model (PDB XXXX), containing CrD in addition to CrH and CrT, aids in understanding the overall positions of the total 123 polypeptide chains that make up the 430 Å repeat of the human cardiac thick filament C-zone and the possible interactions between components. However, we emphasize that the model for CrD should not be taken to represent its detailed atomic structure.

## Data availability

Structural data that support the findings of this study on the structure of the human cardiac thick filament have been deposited in the Electron Microscopy Data Bank. One 2-crown and two 5-crown maps have been deposited, with accession codes EMD-XXXXX, EMD-XXXXX and EMD-XXXXX, respectively. An atomic model of the full 3-crown structure, with myosin heads, tails, titins, and cMyBP-C, has been deposited in wwPDB with accession code XXXX.

## Reporting summary

Further information on research design is available in the Nature Research Reporting Summary linked to this paper.

## Acknowledgments

We thank Drs. Christna Ouch, KangKang Song and Chen Xu for help and training in cryo-EM imaging, Drs. Kyoung Hwan Lee and Greg Hendricks for conventional EM training, and Dr. Niko Grigorieff for use of the Leica EM GP2 plunge freezer. We are grateful to Drs. Tom Irving and Weikang Ma for the unpublished X-ray diffraction pattern of porcine cardiac muscle shown in Extended Data Fig. 3 and Dr. Christine Cremo (University of Nevada) for the gift of gelsolin N-terminal-half plasmid. This work was supported by NIH grants AR072036, HL139883, HL164560, AR081941, HL149164 and HL148785. UCSF ChimeraX, used for molecular graphics, was developed by the Resource for Biocomputing, Visualization, and Informatics at the University of California, San Francisco, with support from NIH grant GM129325 and the Office of Cyber Infrastructure and Computational Biology, National Institute of Allergy and Infectious Diseases.

## Author contributions

D.D. prepared specimens, performed cryo-EM, carried out reconstruction, atomic fitting, refinement, structure analysis, and database deposition. V.N. provided computational expertise, analyzed data. K.S.C. provided curated human heart tissue. R.C. and R.P. carried out analysis of the structure, co-wrote the paper, and obtained funding.

## Additional information

**Correspondence and requests for materials** should be addressed to R.C.

**Reprints and permissions information** is available at www.nature.com/reprints.

**Extended data** follow (ten figures).

**Extended Data Fig. 1.**
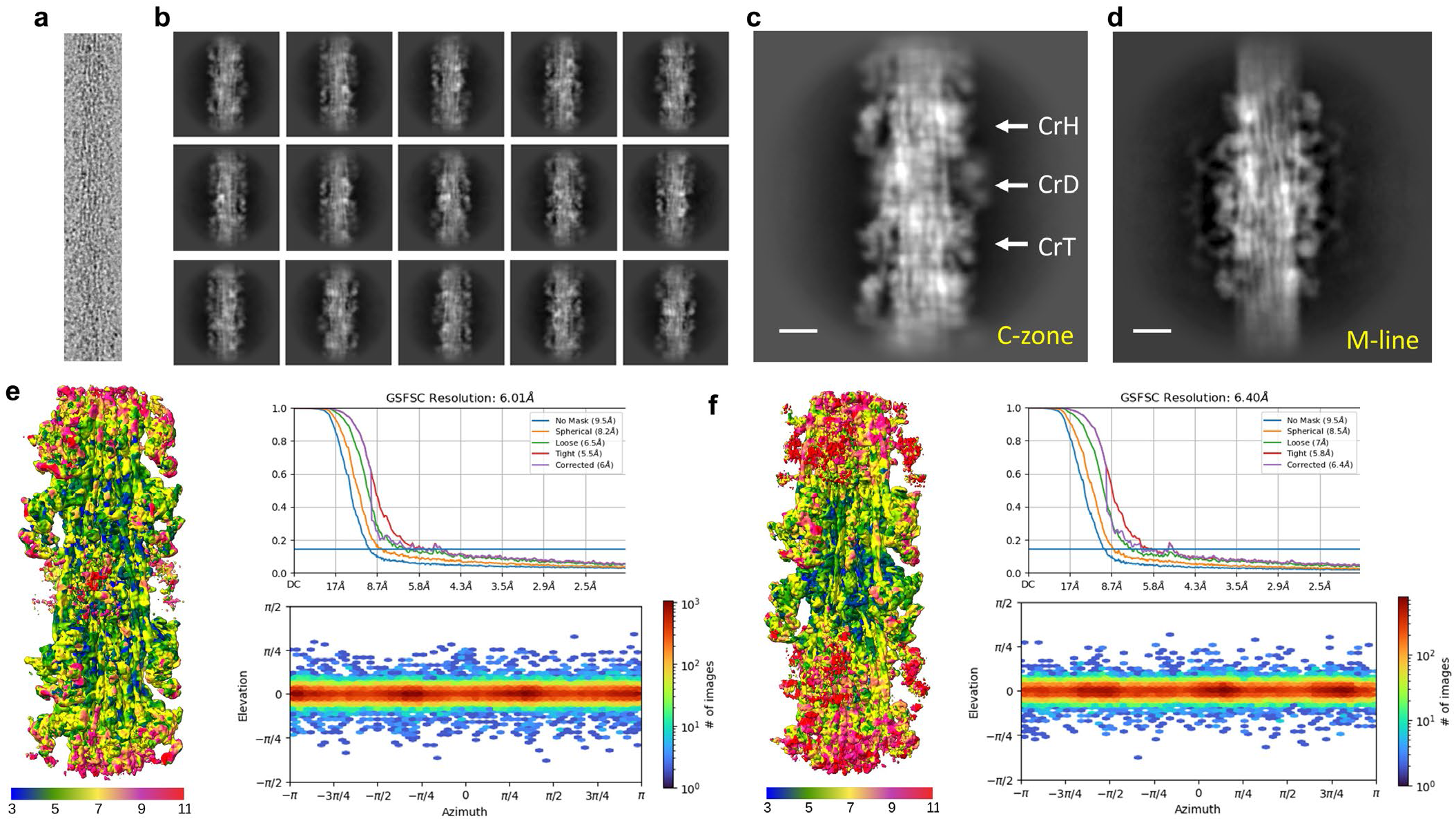
Cryo-EM imaging and data processing. **a,** Cryo-EM image of human cardiac thick filament. **b,** 2D-class averages of 5-crown-long segments of C-zone. **c,** Enlargement of 2D class average. Two IHMs (CrH and CrT) show strong density while the other (CrD) is fuzzy, indicating mobility. **d,** 2D class average of M-line region, included in some images (3D reconstruction to be published separately). **e, f,** The two 5-crown reconstructions produced by CryoSPARC (see Methods), showing local resolution (colored maps), global resolution estimate (gold standard Fourier shell correlation (GSFSC) curves, upper), and heat maps of angles of view of particles used in the reconstruction (lower). The local resolution maps show best resolution in the central 3 crowns; disorder in CrD is indicated by its lower resolution (red) compared with CrH and CrT. Analysis of the maps (see text) shows highest resolution (blue) where interactions occur between components, which would stabilize these regions. The heat maps show a good distribution of angles of view. Most particles are close to in-plane (elevation axis), due to extended, relatively rigid nature of the thick filament, while all rotations about the filament axis are included, with a small preferred-orientation every 120° (darker patches along azimuth axis). Scale bar in **c, d,** 100 Å.

**Extended Data Fig. 2.**
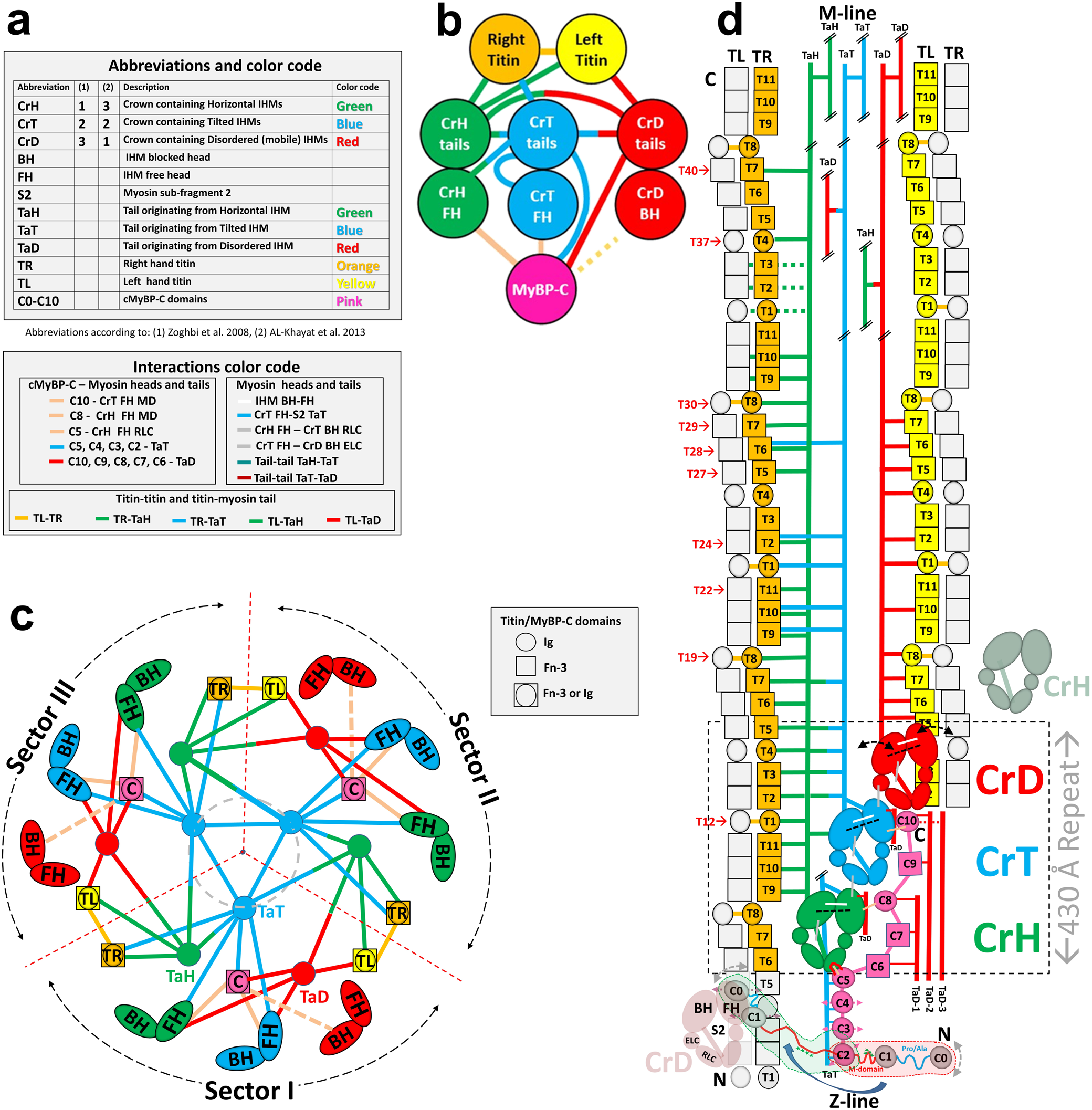
Abbreviations and overview of interactions in 3D reconstruction. **a,** Abbreviations and color coding used to describe reconstruction and interactions in the thick filament. **b,** Interaction network between titin, myosin (heads and tails), and cMyBP-C. (1) titins (TR, TL) interact with each other (at 2 points only), with CrH and CrT tails (TR), and with CrH and CrD tails (TL), but not with cMyBP-C. (2) cMyBP-C interacts with CrH and CrT FHs, and possibly with CrD BH, with CrT and CrD tails, but not with titin. (3) all tails interact with each other (TaH-TaT, TaT-TaD, TaH-TaD) and with themselves (TaH-TaH, TaT-TaT, TaD-TaD). Neither titin, cMyBP-C, nor myosin (heads or tails) hold the thick filament together alone; instead, all orchestrate its interacting, asymmetric structure. **c**. Cartoon showing interactions in the reconstruction viewed transversely, exhibiting three equivalent radial sectors I-III. Due to 3-fold symmetry, the positions of the dividing dotted red lines between sectors are arbitrary. We have chosen them to produce a division in which most interactions occur within a sector, and few between sectors **(ED Fig. 4a)**. The sectors thus defined may correspond to the 3 subfilaments into which vertebrate thick filaments fray at low ionic strength^76^, as these would break the fewest interactions. Only two interactions, between titins TL and TR, in neighboring sectors, would be broken, at T1 and T8. Note how TL and TR pairs from adjoining sectors are nicely accommodated in the space between CrH and CrT IHMs (see also **Fig. 1c, d**). The structure suggests that when filaments are synthesized in the cell, three pre-formed sectors could assemble into a filament when TL and TR strands of different sectors zip together establishing the T1 and T8 interactions **(SI Discussion)**. A central core (gray, dashed circle) contains only CrT tails, interacting with themselves. **d,** Cartoon showing interactions in sector I of the reconstruction, as viewed from the bottom of **c** (i.e. from outside the filament). CrH, CrT and CrD IHMs comprise a 430 Å repeat (dashed rectangle), with adjacent CrD and CrH (pale red and green ghosts) below and above. The dashed black lines in the three colored IHMs denote their tilt: horizontal (CrH) or tilted (CrT, CrD). CrD heads are mobile (curved double-arrows). In the reconstruction, cMyBP-C C5-C10 domains exhibit strong densities, while C2-C4 and C0-M, which are mobile (straight double-arrows), exhibit weaker (C2-C4) or much weaker (C0-M) densities, C0-M possibly docking intermittently on the CrD FH (green dotted fragment). In contrast, in the sarcomere, the mobile C0-M domains (gray, curved double-arrows), may detach from the thick filament, extending perpendicularly and binding to the thin filament (red dotted fragment^44^). Double oblique lines in tails indicate that only partial tails are shown. Red numbers of titin domains refer to the numbering in Fig. 4.

**Extended Data Fig. 3.**
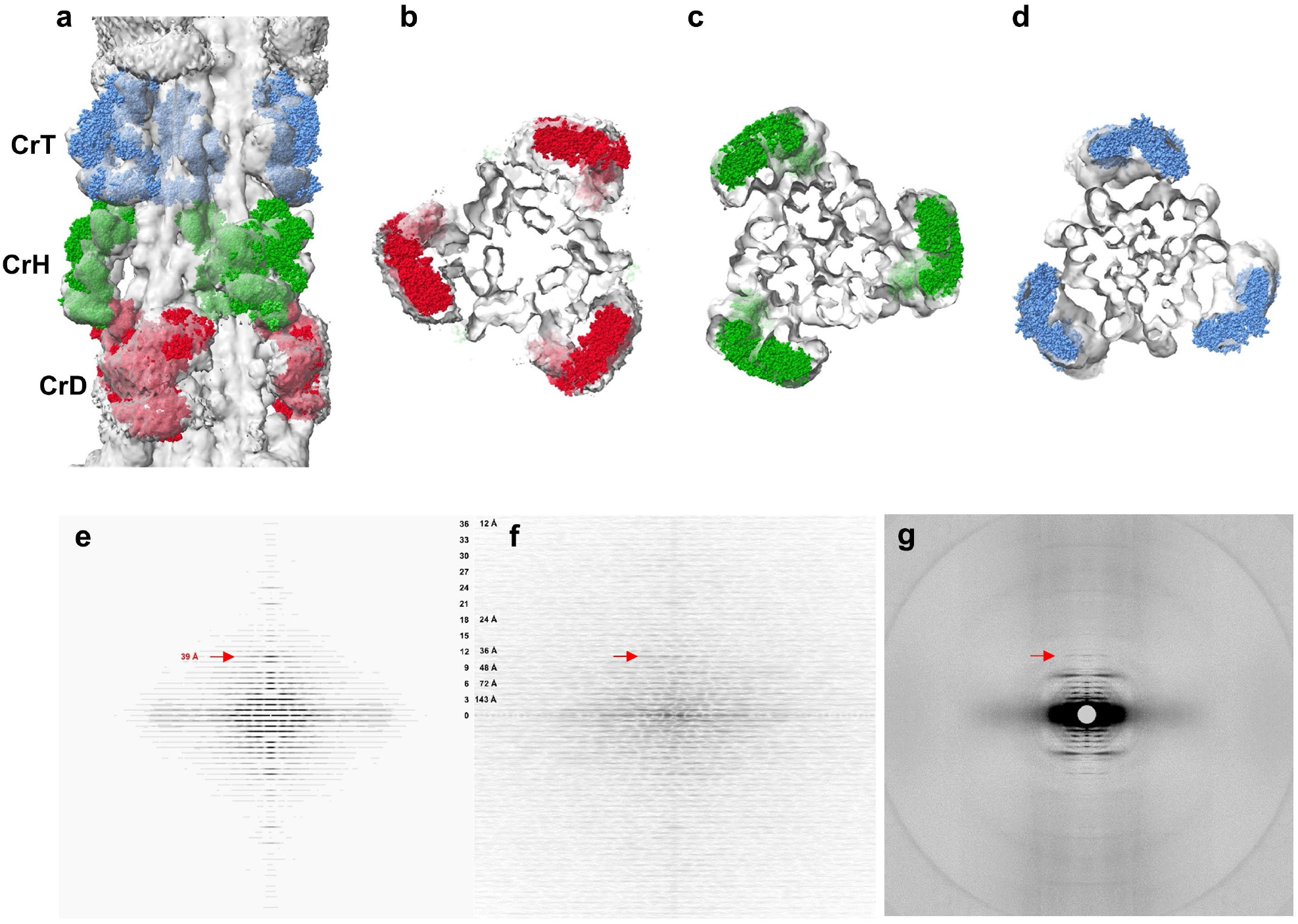
Validation of reconstruction. **a-d,** We compared our reconstruction with X-ray diffraction data, which provide information on filament structure in the lattice of intact muscle. **a,** Longitudinal, and **b-d**, transverse views of our cardiac thick filament reconstruction at the 3 crowns in the 430 Å repeat (gray map), fitted with thick filament atomic model (CrH, green; CrT, blue; CrD, red) based on X-ray diffraction data^22^ combined with previous negative stain reconstructions^18, 19^. The excellent fit of the X-ray model to the cryo-EM map, especially the radial positioning of the heads (center-of-mass ∼135 Å from the filament axis^22^), suggests that the reconstruction is close to the structure in intact muscle, and strongly supports its validity. The previous negative stain reconstructions had suggested a head center-of-mass at ∼95 Å radius (∼40 Å less than the cryo-reconstruction), thought to be due to radial collapse of heads during drying of negative stain^22^—not an issue with frozen-hydrated specimens. **e-f,** We also compared an averaged power spectrum of the reconstruction (rotated at intervals of 10° from 0 to 110°) **(e)** with an averaged power spectrum of selected filaments used in the reconstruction **(f)**. The similarity of the two power spectra, both extending to the 36th order of the 430 Å repeat (12 Å), supports the validity of the reconstruction. **g,** Wide-angle X-ray diffraction pattern of intact, relaxed porcine cardiac muscle at same scale (courtesy of Drs. Tom Irving and Weikang Ma, unpublished data) shows similar myosin layer lines at low angles, further suggesting that our structure is similar to that in intact muscle. A key feature of all patterns is the prominent 39 Å reflection (11th order of 430 Å, red arrows). Previous X-ray studies have suggested that this may be due to titin^77^. Our reconstruction reveals titin unambiguously (see text and **ED Fig. 5**), and we show that the reflection arises from the kinking of its elongated structure, allowing 11 domains to fit into the 430 Å repeat (**ED Fig. 6c**).

**Extended Data Fig. 4.**
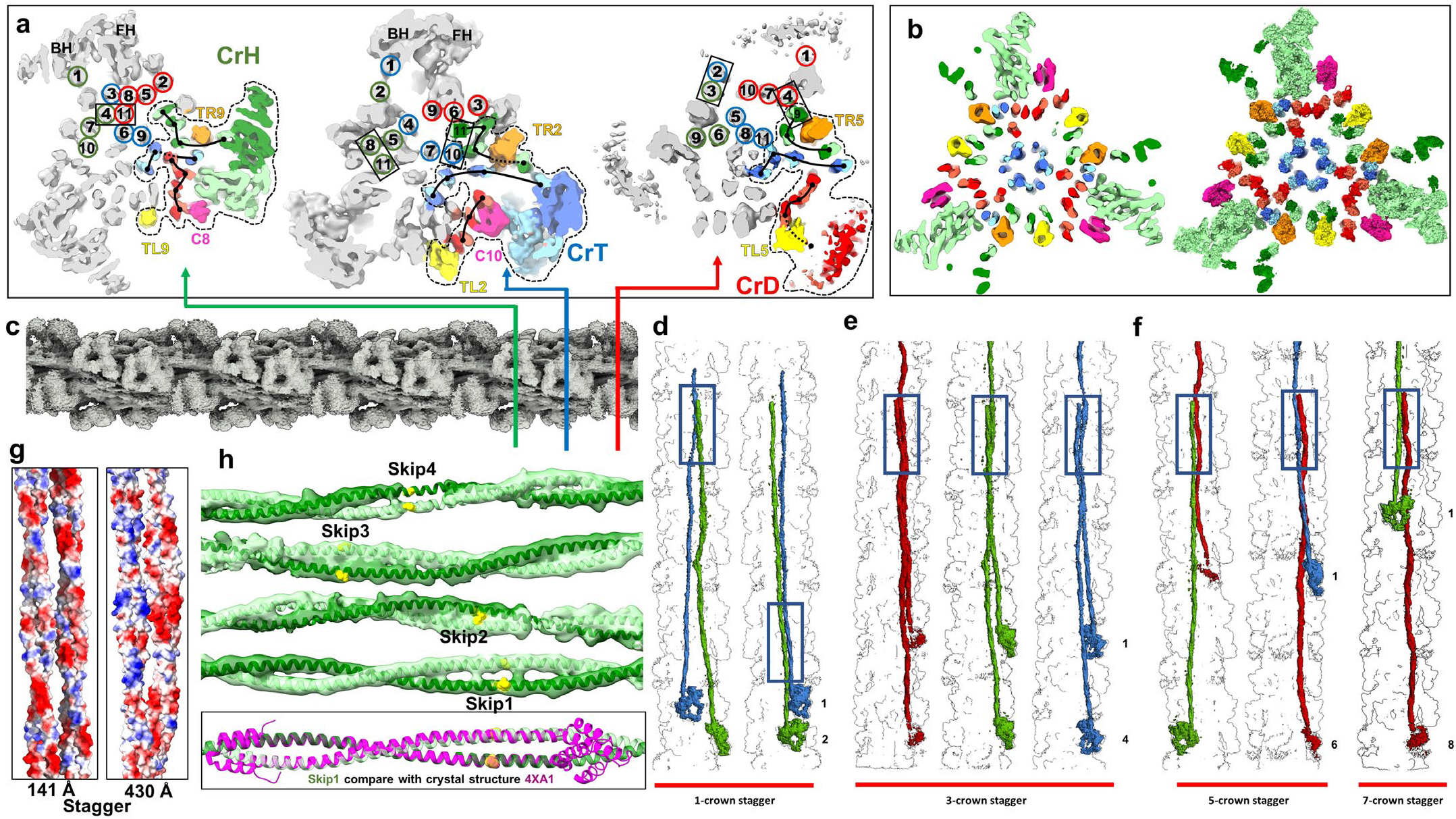
Tracking myosin tails and their interactions. **a,** Transverse slices of reconstruction (looking towards M-line) showing the tail arrangement at crowns H, T, and D, respectively (colored lines in **c**) for TaH (green), TaT (blue) and TaD (red). Tails are tracked by showing their positions at different crown levels (numbers), using the convention devised by Squire^21, 26^, as they travel from their N-terminal origins, at the head-tail junction of each IHM (level 1), to their C-terminal tips (level 11). Black lines in colored sectors replace the numbers, revealing the quite different courses of the 3 tail types as they travel along the filament. For example, TaD lie near the surface while TaT are the most central, forming the filament core. Left and right titins are labeled TL*n* and TR*n*, where *n* is the n^th^ titin domain in the 430 Å repeat. MyBP-C is pink and labeled C*n* (where *n* is the MyBP-C domain^30^). The dotted black lines show one sector (colored) at each crown. Rectangles show examples of interactions of tails staggered by 1, 3, 5, and 7 crowns. Note sheets of CrD tails (red) staggered by 3 crowns (430 Å) near surface of backbone, forming a binding platform for cMyBP-C (see text). **b,** Cross-section at CrH, showing map (left) and corresponding space-filling atomic model (right), revealing clear examples of contact (thus interaction) between myosin tails with various staggers. Similar contacts are seen at other levels. **c,** longitudinal view of map (M-line at right) showing where slices in **a** are cut. **d-f,** Longitudinal views of reconstruction showing interactions of tails staggered by 1, 3, 5 or 7 crowns (∼ 143, 430, 715 or 1,001 Å), as predicted by^23^, corresponding to differences in tail numbers of 1, 3, 5, and 7 in **a**. Main sites of contact are inside the rectangular boxes. The most common sites are the distal LMM regions of each tail, including the ACD at the C-terminus (top boxes), consistent with its requirement for filament assembly (see text). The most prevalent stagger (3-crowns, 430 Å) is confined to homologous tails (TaH-TaH, TaT-TaT, TaD-TaD) **(e)**. Heterologous tail combinations are staggered by 5 crowns (TaT-TaD, TaD-TaH) and one (TaH-TaD) by 7 crowns **(f)**. One heterologous pair (TaH-TaT) exhibits a 141 Å stagger between S2s in the same sector **(d, right)** and between LMMs (including the ACD) from different sectors **(d, left). g,** Examples of charge interactions between S2’s staggered by 141 Å and LMMs by 430 Å. **h,** Map and model at the 4 skip residues in TaH tails **(SI Table 1b)** and comparison of skip 1 in the filament (green) with corresponding crystal structure of skip 1 (PDB 4xa1, pink^24^). The similarity gives support for the relevance of this X-ray model to the native structure.

**Extended Data Fig. 5.**
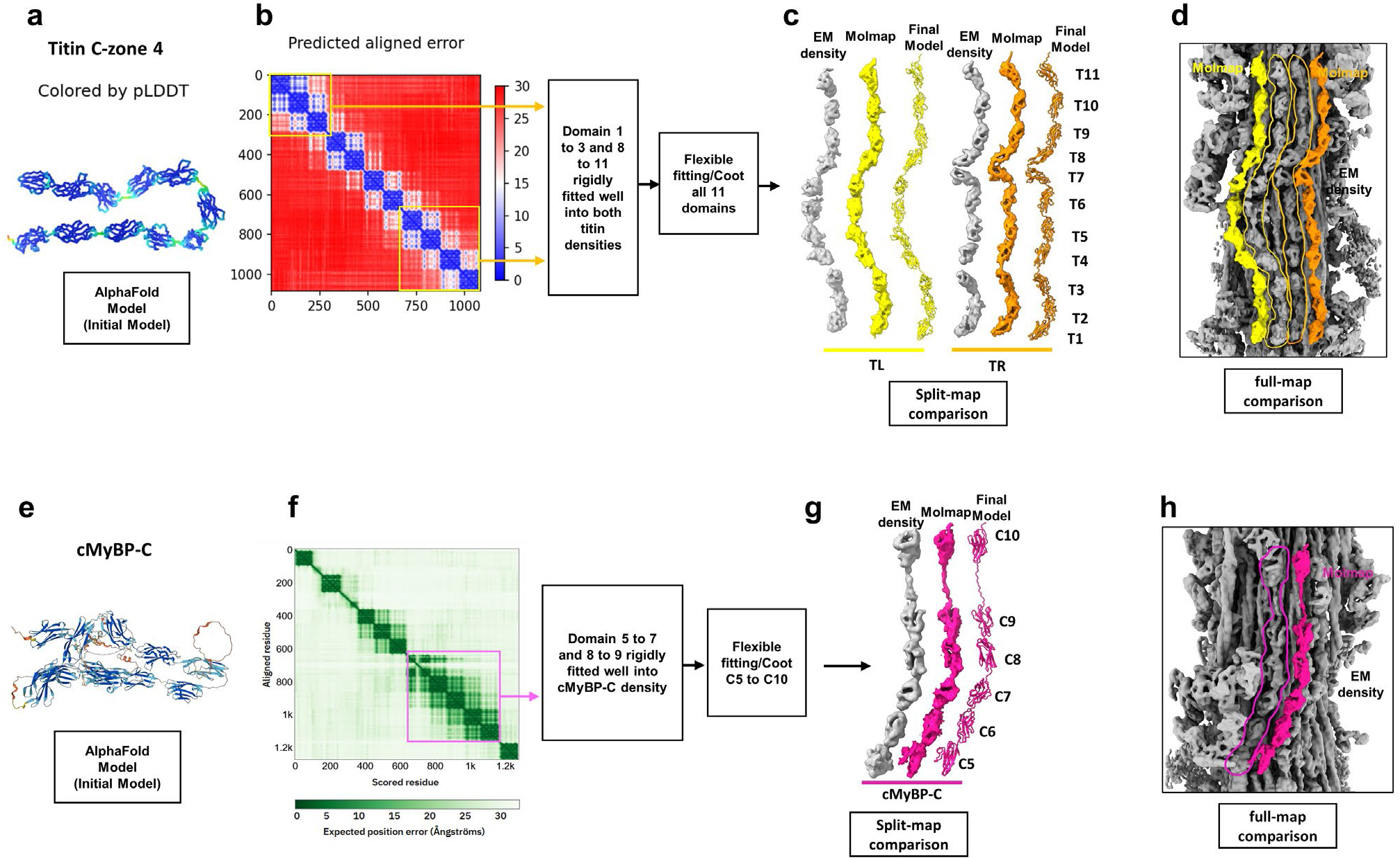
Domain identification and model building of titins and cMyBP-C. **a, e.** AlphaFold^34^-predicted models of the two titins **(a)** and cMyBP-C **(e)**. The sequence for titin super-repeat 4 of the C-zone was used, as each super-repeat is unique. **b, f,** Domains of titin and cMyBP-C showed high confidence. pLDDT (predicted local distance difference test) was ∼90 (high confidence; blue in **a)**. Predicted aligned error (PAE) was also good (< 5 Å). From PAE plots, portions with inter-domain confident regions (yellow and pink boxes in **b** and **f)** were initially rigid-body fitted to the reconstruction (‘fit in map’ in ChimeraX), and other domains then connected using Coot and flexible fitting (MDFF, NAMDINATOR) to build the final atomic models. **c, g,** Atomic models of TL, TR and cMyBP-C. Maps (yellow, orange, pink surfaces) were generated from the models, using Molmap in ChimeraX, to compare with the EM density in the reconstruction (gray surface maps segmented from full reconstruction). The predicted maps showed excellent agreement with the actual map for every domain, giving confidence in our atomic models for the two titins and cMyBP-C. An especially striking prediction of AlphaFold was the long linker between cMyBP-C domains C9 and C10 **(g)**, which fitted precisely into our EM map. **d, h**, Predicted density maps (yellow, orange, pink), based on atomic models **(c, g)**, placed on the full reconstruction (gray), for comparison in the context of the entire map (actual densities outlined by yellow, orange and pink borders). The similarity of the actual and generated maps confirms our confidence in the assignment of cMyBP-C and titin domains and their near-atomic structures.

**Extended Data Fig. 6.**
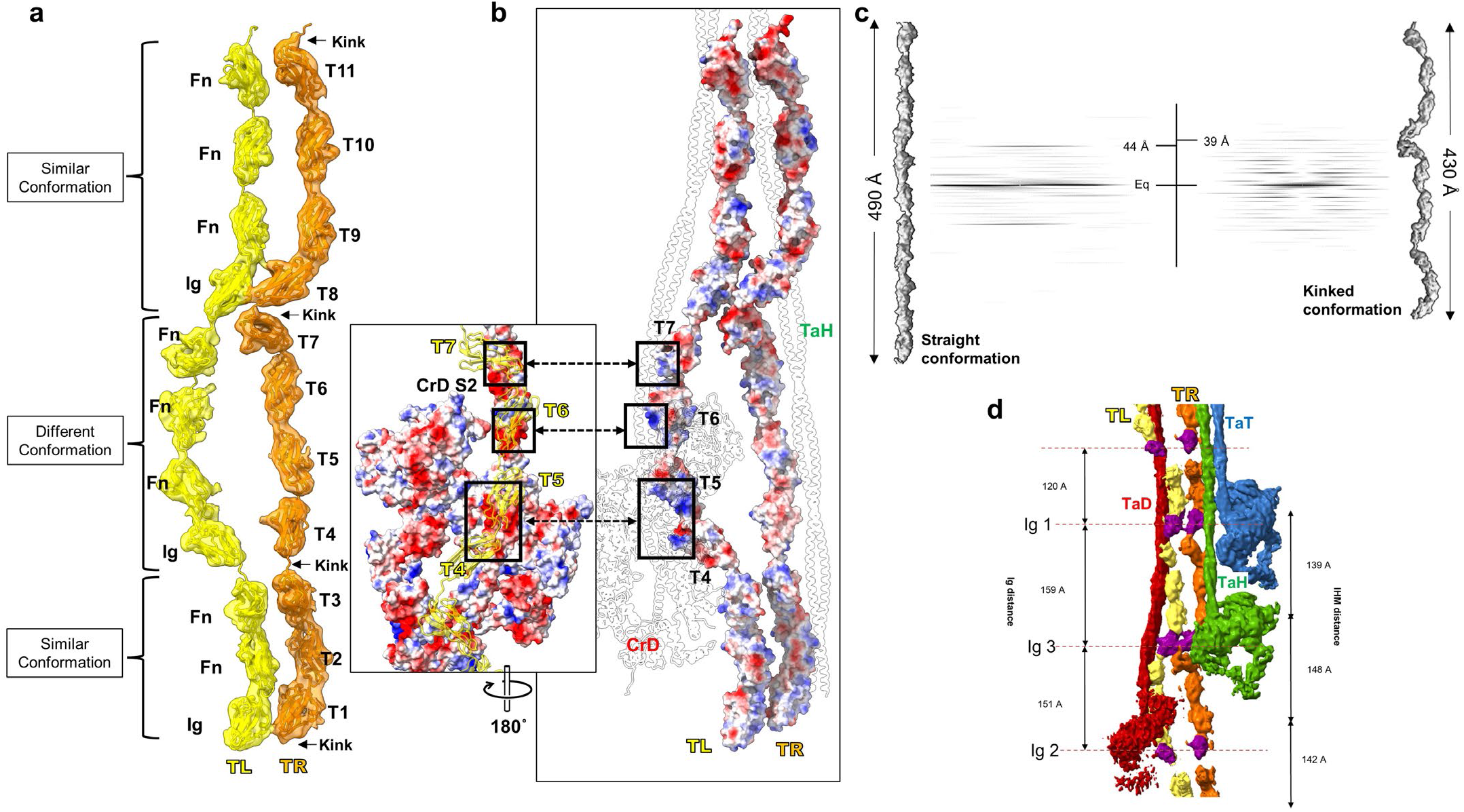
Different conformations of titins TL and TR. **a,** The two titin densities (TL, TR) in one 430-Å super-repeat of a sector contain 11 Ig and Fn domains, and their overall conformation is not straight. The titin maps segmented from the reconstruction show that, due to the presence of 3 kinks (arrows), and a slight curve in the T1-T3 and T8-T11 regions, one super-repeat (11 domains) fits precisely into 430 Å, matching exactly the triplet repeat of myosin heads (CrH-CrT-CrD). The kinks, all occurring at an Ig-Fn junction, demarcate three parts of the super-repeat: 1. T1-T3 (Ig-Fn-Fn); 2. T4-T7 (Ig-Fn-Fn-Fn); and 3. T8-T11 (Ig-Fn-Fn-Fn). Parts 1 and 3 of TL and TR have similar conformations, and are approximately parallel to each other. However, the conformation of the middle section (Part 2) differs between TL and TR. In TL, T4-T7 is rotated ∼90° with respect to TR, positioning it at a higher radius above the surface of the backbone, where it becomes available for interaction with CrD S2 as shown in **b. b,** Because of this conformational difference, the middle portions of TL and TR interact differently with different tails. The bent and raised conformation of T5-T6-T7 in TL binds to proximal S2 of CrD with mostly electrostatic attraction (inset), while the straight conformation of T5-T6 in TR binds to distal S2 of CrH **(ED Fig. 7)**, generating an axial shift of 148 Å between CrH and CrD (see **d**). **c,** Previous studies speculated that the 39 Å reflection in the X-ray diffraction pattern of relaxed muscle **(ED Fig. 3g)** may be due to titin^77^. Here we demonstrate this directly. Fast Fourier transformation (FFT) of the titin strand, segmented from the reconstruction in its native (kinked) conformation (right), produces a meridional reflection at 39 Å, the 11^th^ order of the 430 Å repeat (seen as layer lines at lower angles). If titin is computationally straightened (left), the reflection moves towards the origin (∼44 Å), producing a reflection that is not observed in the relaxed X-ray pattern. Agreement of the reconstruction with X-ray data from relaxed intact muscle **(ED Fig. 3g *vs.* e, f)** implies that titin is kinked in the native state, allowing the 11 domains of its super-repeat to fit into 430 Å. The FFTs were computed from 8 titin super-repeats of the kind shown in **c**, laid end-to-end, to create a strand similar to that in one C-zone of the thick filament. See also **ED Fig. 10. d,** Past studies suggested that the uneven spacing of titin’s 3 Ig domains in the super-repeat might be responsible for the uneven spacing of the 3 myosin crowns in the 430 Å repeat. Our reconstruction enables us to test this idea. We find an approximate correlation between the positions of the 3 Ig domains (purple) and the motor domains of the 3 IHMs. Ig1 (T1) correlates with CrT, Ig2 (T4) with CrD, and Ig3 (T8) with CrH. But we see no precise correlation that would suggest that the 3 Ig domains directly position the crowns (see crown and Ig spacings on figure). This is not surprising, given the absence of any direct titin-head interaction in the structure. Instead, titin positions the crowns through interaction of TL and TR with CrH and CrD tails, while CrT is positioned by interaction of TaT with TaH (see text).

**Extended Data Fig. 7.**
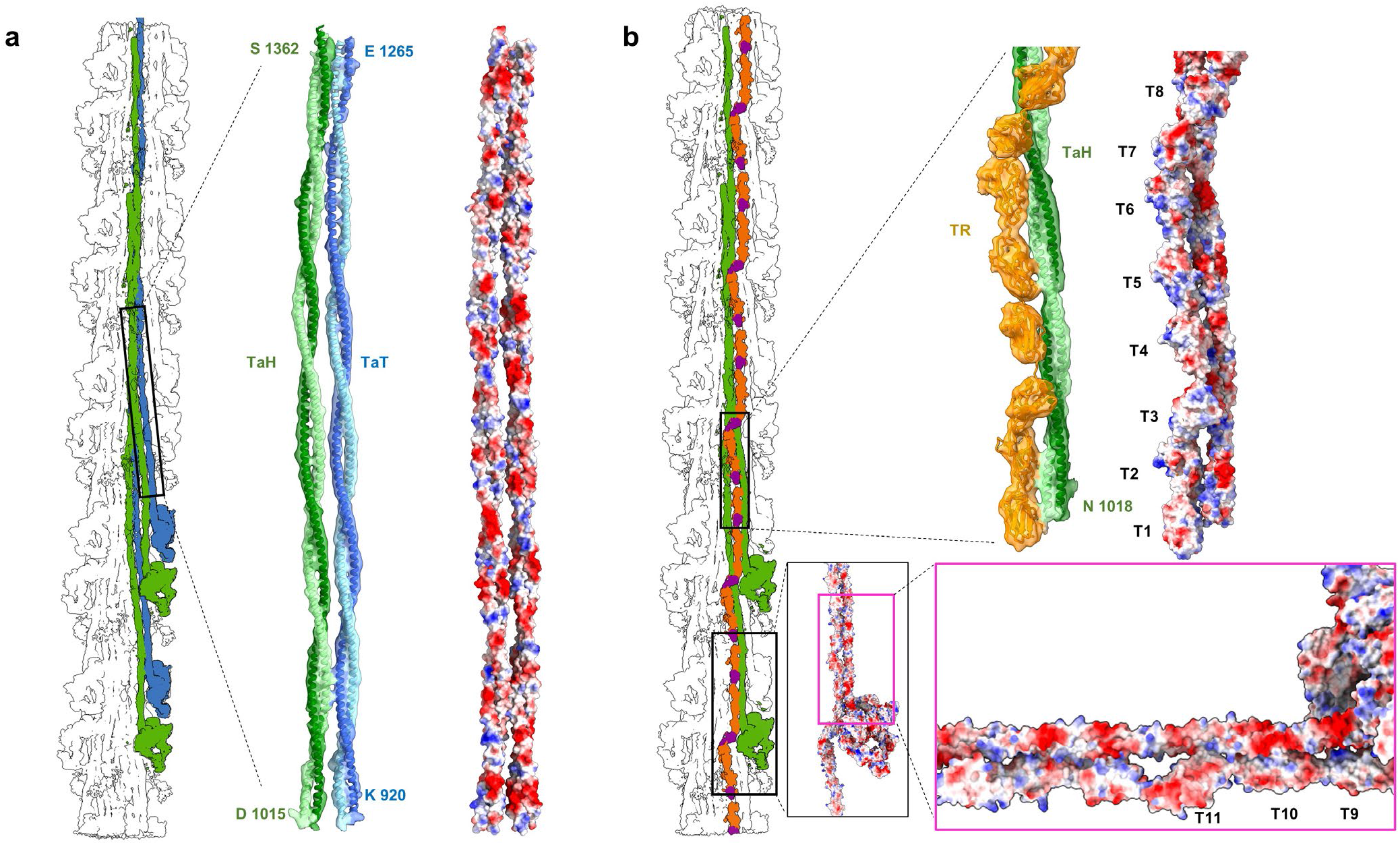
Titin-tail and tail-tail electrostatic interactions create the unique 3-crown repeat of myosin molecules in the C-zone. Myosin tails form extensive, mostly electrostatic, interactions with each other and with titin, which combined determine the axial and azimuthal positions of CrH, CrT, and CrD. **a, b** show representative examples. **a,** TaH and TaT (atomic models fitted into density map) run together in the S2 region (black rectangle) **(Fig. 3)**, forming charge-charge interactions (right: red, negative; blue, positive), which shift CrT ∼141 Å axially with respect to CrH (*cf.*^23^; see also **ED Fig. 4**). **b,** CrH tails also form electrostatic interactions with TR, involving all 11 domains of the 430 Å titin super-repeat **(Fig. 4c).** Interactions involve both proximal and distal S2 (bottom and top right, respectively), whose ∼42 Å charge repeat^23^ matches the ∼42 Å spacing of the charged-surface titin domains. This is the most extensive titin-tail interaction in the reconstruction and is responsible for placing CrH crowns 430 Å apart, by matching them to the 430 Å length of the TR 11-domain super-repeat in its kinked conformation on the filament surface **(ED Fig. 6c)**. In summary, TaH-TR and TaH-TaT interactions (shown here), and TaD-TL interaction (**ED Fig. 6b**), are the driving force for organizing the three crowns (CrH-CrT-CrD) in a quasi-helical arrangement in the human cardiac thick filament in the relaxed state.

**Extended Data Fig. 8.**
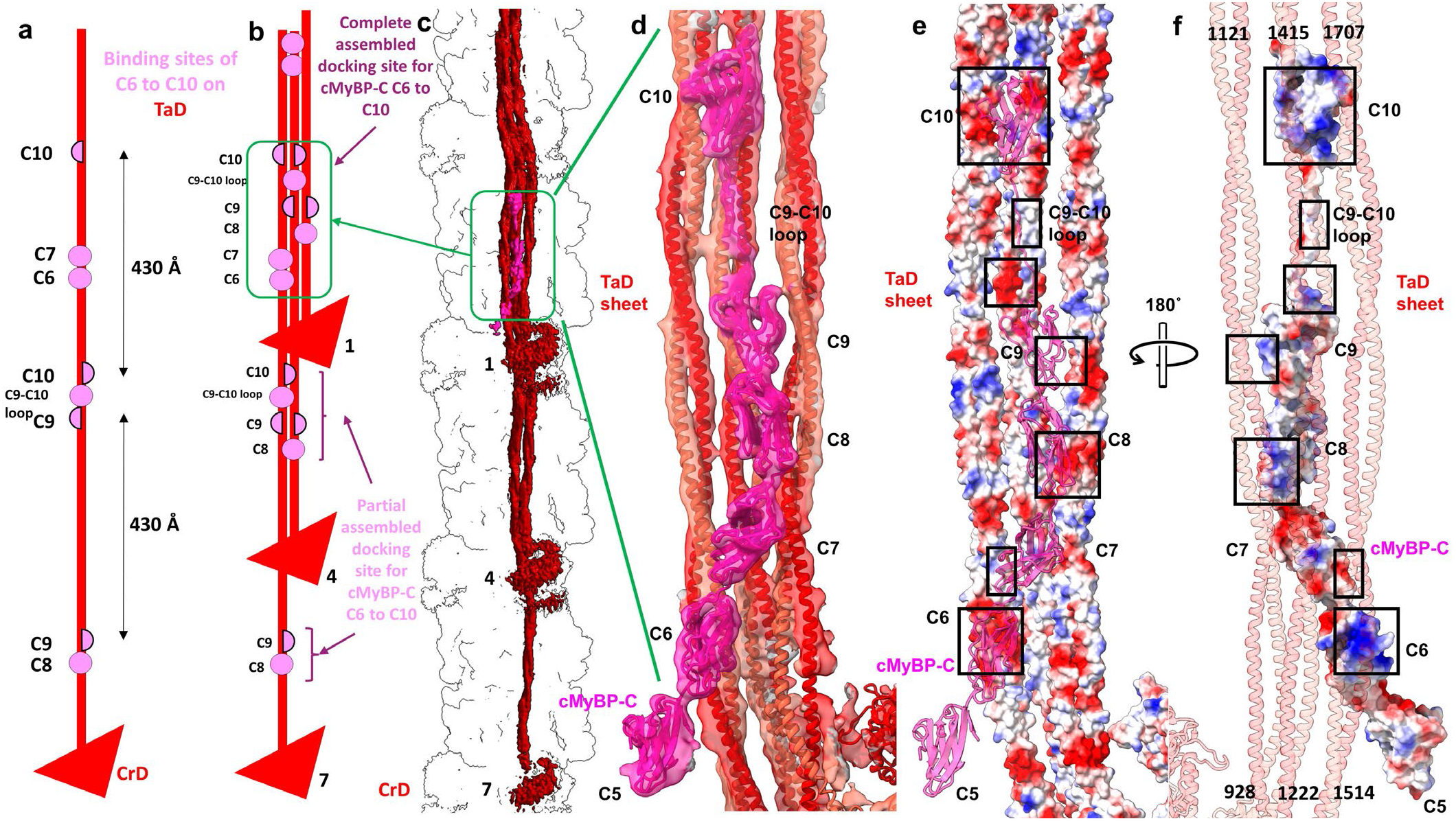
cMyBP-C position is determined by binding to a specific coalescence of CrD tails. The reconstruction shows that C-terminal domains C6 to C10 of cMyBP-C dock onto the filament backbone through interaction with only one type of tail (CrD tails, TaD). There is no interaction with titin, although titin is indirectly involved by positioning the tails. The complete cMyBP-C binding site is formed when three CrD tails, from three consecutive 430 Å repeats, lie side-by-side, creating a sheet-like docking platform that can make complementary charge interactions with the cMyBP-C domains. **a,** Distribution of C6-C10 binding sites at different points along a CrD tail. Complete/partial binding sites are shown by pink circles/semicircles. **b,** When CrD tails from 3 levels assemble to form the docking platform, a complete binding site is created (green box). C6 and C7 interact with level 7 TaD LMM; C8 with level 1 S2; C9 with level 1 S2 and level 4 LMM; C9-C10 linker with level 4 LMM; and C10 with level 4 and level 7 LMMs. The docking of C6 to C10 on the backbone thus requires a particular arrangement of CrD tails, from levels 1, 4 and 7, 430 Å apart. Based on titin’s specific 11-domain super-repeat in the C-zone (determining myosin tail locations), and quite different 7-domain super-repeat in the D-zone^27^, this specific arrangement of CrD tails occurs only in the C-zone. This may explain cMyBP-C’s confinement only to this region of the thick filament. In addition to tail binding, C8 and C10 (both Ig domains) interact with the FH motor domain of CrH and CrT **(Fig. 4)**. **c, d,** Atomic model fitted to map shows details of interactions in the complete cMyBP-C binding site on the 3 CrD tails. **e, f,** Surface charge depiction of tails **(e)** and cMyBP-C, rotated 180° **(f)**, suggests that binding occurs mainly through electrostatic attraction (boxes show complementary charges).

**Extended Data Fig. 9.**
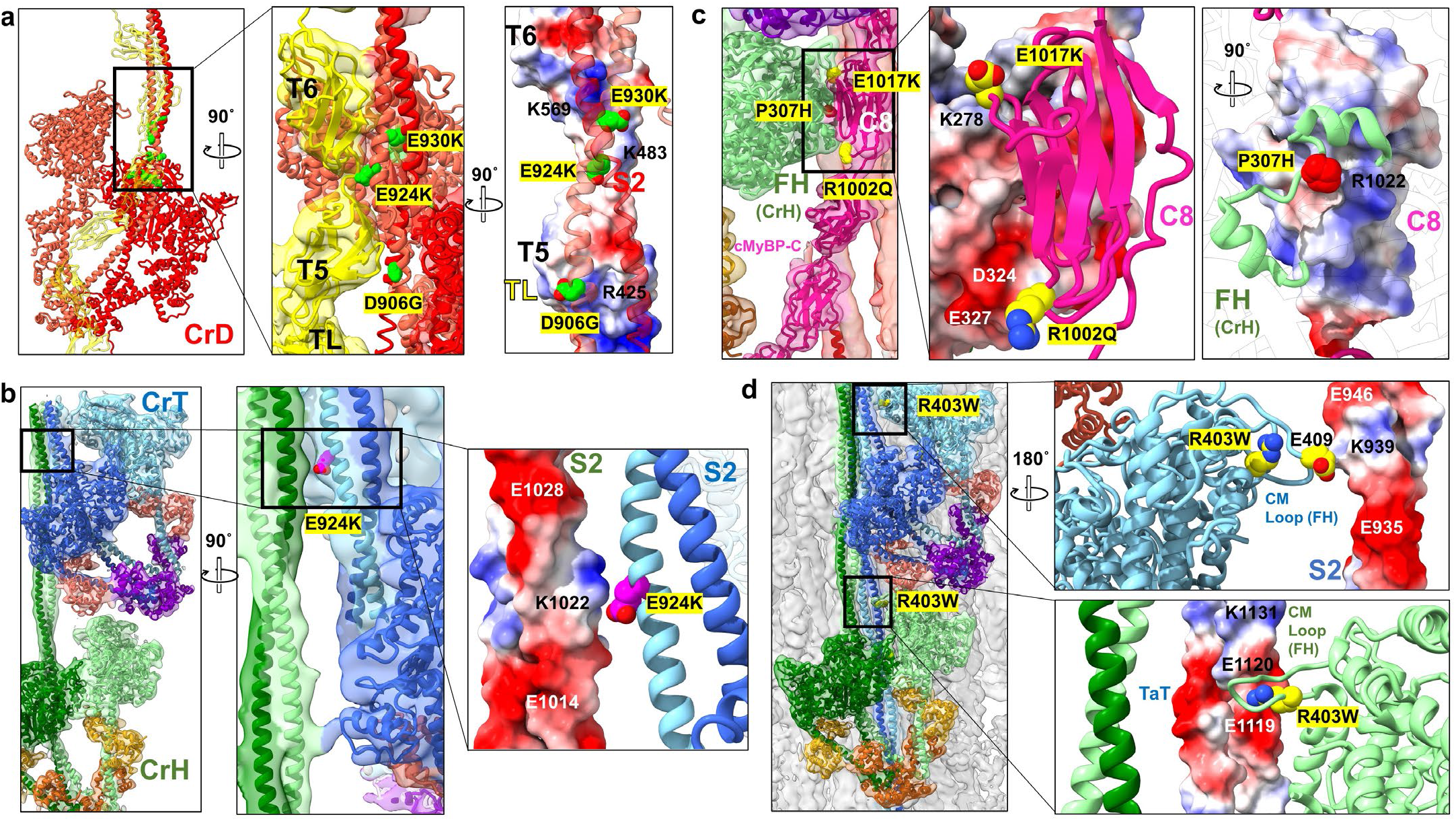
Examples of mutations clustering at intermolecular interaction sites, whose location suggests novel mechanisms of HCM pathogenicity. The map reveals that many HCM pathogenic mutations in cMyBP-C, and in myosin tails and heads of specific crowns (CrH, CrT and CrD), are in intermolecular interfaces in the native filament. These mutations could not be mapped previously due to use of the tarantula filament IHM, lacking cMyBP-C and titin^5, 6^. **a**, **Mutations affecting the tail-titin interface.** Mutations in ring 1 (D906G) and 2 (E924K, E930K) of CrD S2^53^ occur at the site of interaction with TL domains T5 and T6. Surface charge depiction showing positively charged patches on TL (blue on right inset) suggests that these mutations, involving loss or reversal of negative charge, would weaken binding of CrD S2 to TL. This could interfere with transmission of tension by titin to the cMyBP-C–TaD binding site, and thus disrupt possible mechanical signaling mechanisms (see text and **ED Fig. 10**). **b, Mutations affecting tail-tail interfaces.** Mutation E924K in **(a)** is also involved in a tail-tail interaction, in this case not CrD, but the S2s of CrH and CrT. E924K charge reversal on CrT S2 (right inset) would be expected to weaken interaction with positive charge (blue) on CrH S2, thus impairing stability of the tail network. Disruption of function in these 2 different ways (tail-TL and tail-tail) could make the E924K mutation especially pathogenic. **c**, **Mutations in cMyBP-C and the myosin MD affecting the cMyBP-C–myosin head interface.** Pathogenic mutations in the myosin head (P307H) and the cMyBP-C C8 domain (R1002Q, E1017K) both occur in the interaction interface of C8 with the FH MD of CrH. Surface charge depiction suggests that charge loss or reversal on C8 (left inset) and on CrH FH (right inset) would weaken this interface, impairing cMyBP-C’s stabilization of the CrH FH. This could disturb its SRX state and interfere with proposed mechanical signaling mechanisms **(ED Fig. 10). d, Mutations in the myosin MD affecting multiple interfaces with tails.** The classic mutation R403W/G/L/Q, in the CM loop of the MD, has been widely studied but not fully explained^78^. The reconstruction suggests that it could have multiple effects, by impairing head-tail interactions differently in CrH and CrT FHs. In the upper inset, the CrT CM loop interacts with its own S2, while in the lower, the CrH CM loop interacts with S2 from the CrT below. Loss of positive charge in the former may strengthen binding to the positive charge K939, stabilizing the IHM, while in the latter it would weaken interaction with the CrT tail (E1119/E1120), again affecting mechanical signaling mechanisms **(ED Fig. 10)**. In summary, due to the various environments of myosins in a single 430-Å repeat, a single mutation can affect multiple interactions, with different pathogenic consequences.

**Extended Data Fig. 10.**
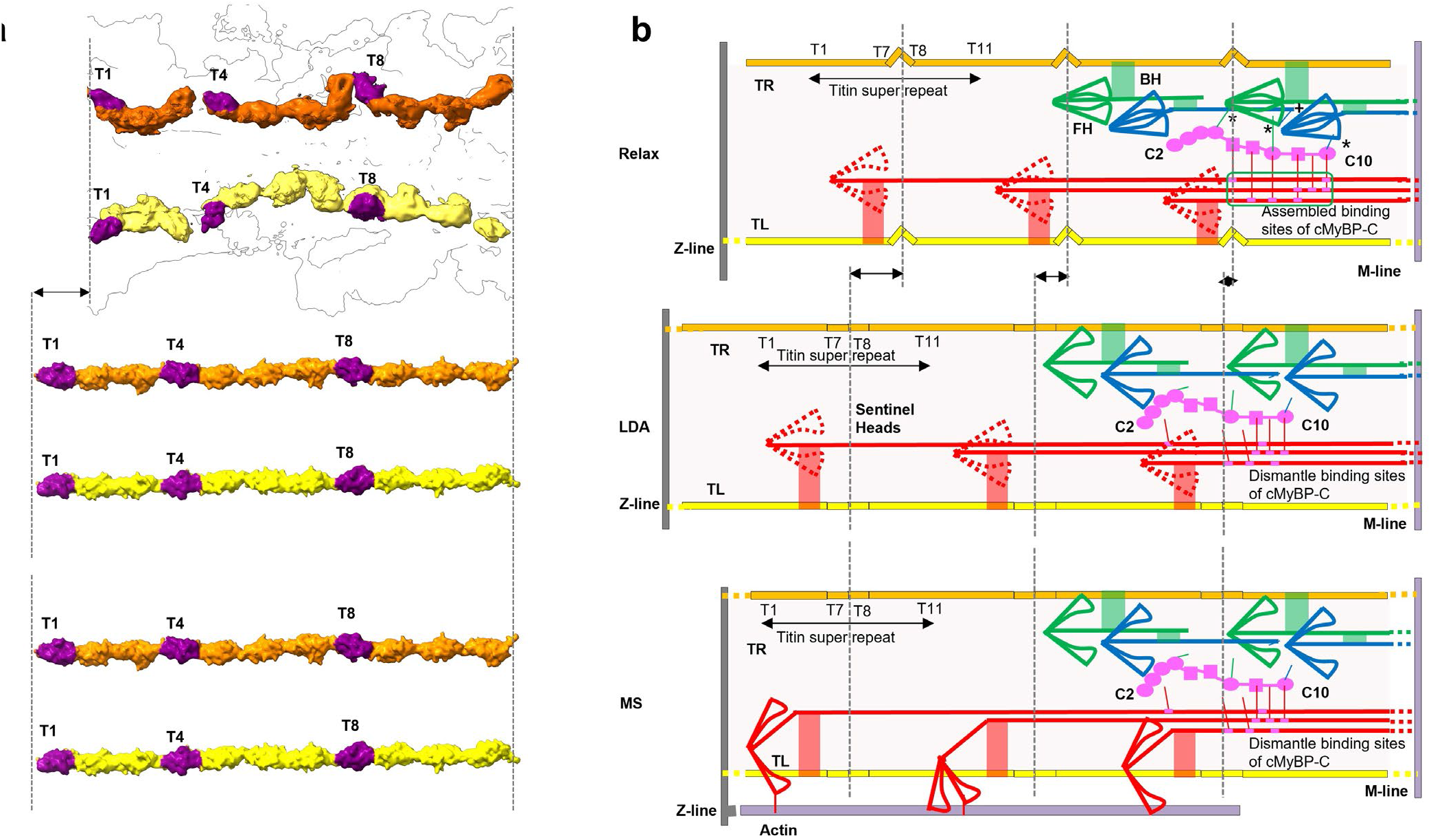
Proposed mechanism of length-dependent activation (LDA) and mechanosensing (MS) in cardiac thick filament involving titin, cMyBP-C and myosin tails. Our reconstruction reveals connections of titin to myosin and myosin to cMyBP-C, that may underlie LDA and MS. **a, b. Upper (relaxed):** In relaxed state (right), TL and TR are kinked as in our reconstruction (left). Myosin heads form IHMs; cMyBP-C binds to docking site on CrD tails (green box; **ED Fig. 8**) and to CrH and CrT heads (asterisks) **(Fig. 5c-f)**; and TL and TR interact with myosin tails (red, green connections). In addition, CrH FH CM loop binds to TaT tail (“+” in **b** (upper); Fig. 6g**, h**)**. Middle (LDA):** Elongation of the sarcomere at end-diastole (right) stretches TL and TR, reducing the kinks (left), supported by an increase in the spacing of the 39 Å X-ray reflection upon stretch^79^; see **ED Figs. 3g**, 6c). Tension-induced translation of titin domains pulls on connected myosin tails, dismantling cMyBP-C binding sites on CrD tails and on CrH and CrT heads. Loss of the stabilizing influence of cMyBP-C on CrH and CrT, augmented by weakening of the CrH FH CM loop-TaT interaction (“+” in **b,** upper) under tension, releases heads for actin interaction, accounting for progressive enhanced force of systole that follows corresponding sarcomere length increase (= LDA). **Lower (MS):** In mechanosensing (right), CrD heads that performed a “sentinel” role for detecting thin filament activation^80, 81^ in relaxed and LDA states, are now active and produce tension by interacting with actin (right). This stretches titin in a similar way to LDA (left), with similar consequences, enhancing contractility. In addition to the above cMyBP-C interactions, there are also CrD tail interactions with CrH FH and CrT FH, which could stabilize these IHMs **(Fig. 6j, k, m)**. These interactions would also be broken upon sliding of CrD tails, contributing to LDA and MS.

## SUPPLEMENTARY INFORMATION

### Supplementary discussion

Our structure suggests a plausible model for thick filament assembly in vivo. Titin is laid down first^27^ and recruits myosin II into nascent filaments. S2 of a myosin molecule binds to TR at each of the 11 domains in its 430 Å super-repeat, supplemented by additional, more distal, interactions with TR, forming CrH crowns **(Fig. 4c)**. Additional CrH myosins bind to other 430 Å super-repeats of TR. CrT myosin S2s interact with CrH S2s, staggered by ∼141 Å **(Fig. 3e)**^23^, and also interact with a short, nearby region of TR **(Fig. 4c)**; their distal tails form the filament core. More CrT’s add on, creating their own 430 Å repeat. TR is now fully occupied with bound myosins **(Fig. 4c, e)**, requiring a second titin (TL) to bind CrD tails, which occurs on the surface of the filament due to the different conformation of TL from TR. As myosins find their positions by binding to titin, their tails also bind to each other through the stagger interactions shown in **Fig. 3c and ED Fig. 4d-f**, consolidating the backbone. Other orders of incorporation of myosin molecules are possible, or simultaneous addition might occur. cMyBP-C is recruited last^82^, only after the sheet of 3 CrD tails is formed to provide its docking platform on the filament backbone **(ED Fig. 8)**; we see no evidence for direct binding of cMyBP-C to titin. Individual sectors may form in this way first, zipping together via TL-TR Ig domains at T1 and T8 **(ED Fig. 6a, b)** and tail-tail interactions between sectors, to form the mature, 3-fold symmetric filament. The mature filament has a relatively open backbone structure **(Figs. 1e, 2b; ED Fig. 4a, b)**, which we suggest would explain the ability of myosin molecules to dynamically exchange into and out of the filament during protein turnover^83^.

